# Evaluating the Generalizability of Machine-Learning-Guided Directed Evolution Across Protein Fitness Landscapes

**DOI:** 10.64898/2026.11.24.769005

**Authors:** Rajarshi SinhaRoy, Alexander Tauber, Georg Künze

## Abstract

Machine-learning-guided directed evolution (MLDE) prioritizes promising protein variants for experimental testing, but performance depends on campaign design. We present MLDEx, a systematic benchmark of 557,143 simulated campaigns across 16 protein fitness landscapes, covering 28 representations, eight regressors, three acquisition rules, four initial training-set sizes, and four batch sizes. Going beyond previous benchmarks, MLDEx jointly evaluates these design choices and tests whether recommended configurations generalize to unseen protein landscapes. With small initial sets, ML guidance required 2.2–2.8 times fewer assays than random screening for comparable success, but this advantage diminished with larger sets. At 200 total assays, zero-shot screening achieved 56.2% success versus 44.6% for MLDE, with similar performance at larger budgets. Representation–regressor rankings varied substantially between landscapes: configurations selected on 15 landscapes performed 13.3 percentage points worse on the held-out landscape. MLDEx identifies broadly applicable campaign-design principles but cautions against transferring model rankings to new proteins.

**TOC Figure:** 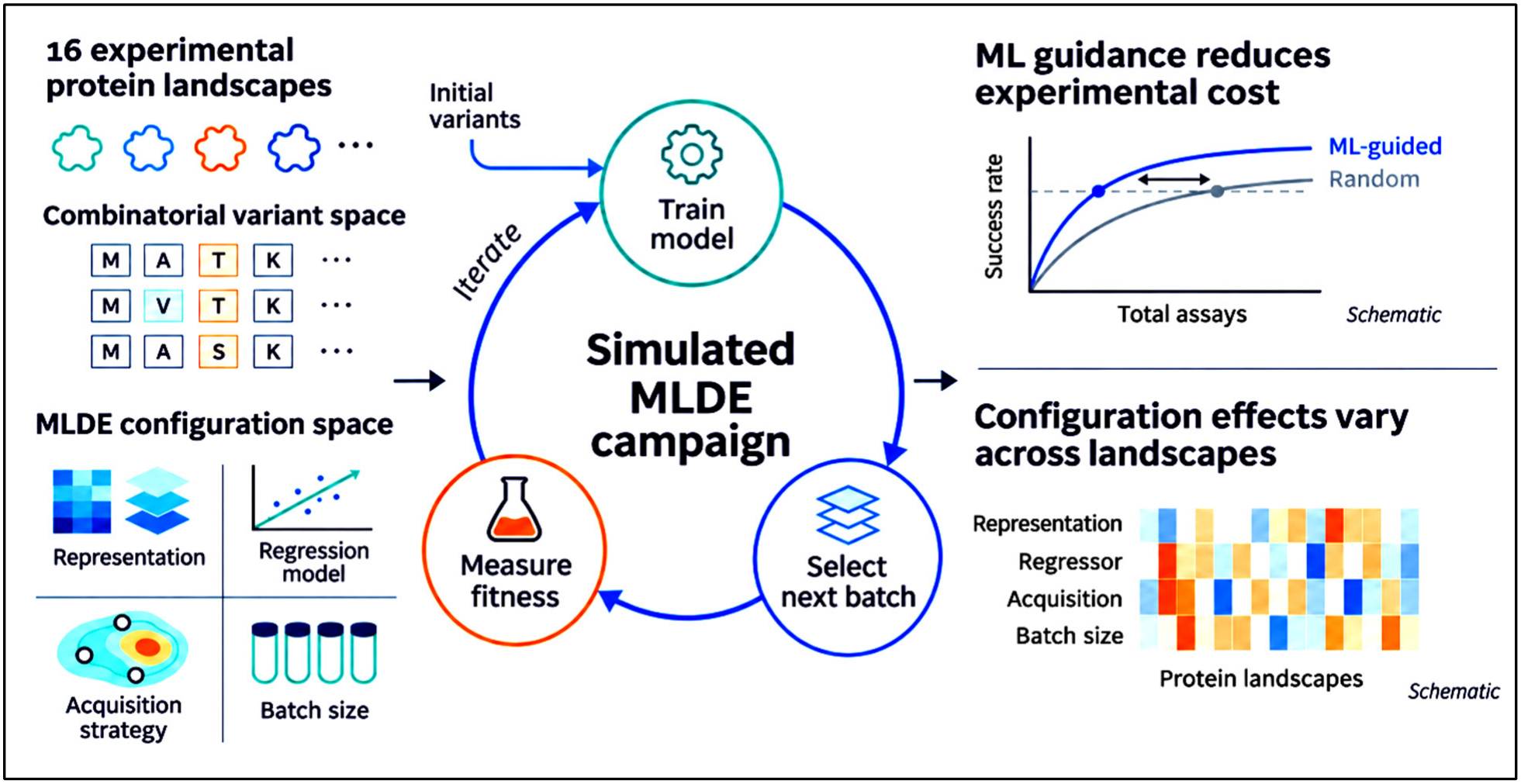

## INTRODUCTION

Directed evolution has become a powerful approach for improving protein function through iterative cycles of sequence diversification and experimental screening.^1^ However, the vast protein sequence space makes exhaustive experimental characterization impractical, particularly for combinatorial libraries in which even a few variable positions can generate millions of possible variants.^1^ Machine-learning-guided directed evolution (MLDE) addresses this challenge by using experimentally characterized variants to train a predictive model that prioritizes candidates for subsequent rounds of screening.^2,3^ By directing experimental measurements toward variants predicted to have high fitness, MLDE can potentially reduce the number of assays required to identify improved variants.

The performance of an MLDE campaign depends on several design choices: the representation used to encode protein variants, the regression model used to predict fitness, the acquisition rule used to select variants for the next round of screening, the size of the initial training set, and the number of variants assayed per round. Individual studies have examined these choices and reported strategies that improve MLDE performance in particular experimental settings.^3^ However, the relative performance of different approaches may depend on the underlying protein fitness landscape. This issue is well recognized in machine learning, where the ranking of competing methods can vary substantially across benchmark datasets and tasks.^4,5^ Whether the relative performance of MLDE configurations shows similar landscape dependence has not been systematically evaluated. Because these choices must often be made before the fitness landscape is characterized, recommendations are most valuable if they generalize to previously unseen landscapes.

Previous work can be divided into three groups. The first comprises standardized benchmarks for variant-effect prediction. Dallago et al. assembled FLIP, a collection of fitness-landscape tasks with defined training and test splits for comparing protein representations.^6^ Notin et al. extended this approach in ProteinGym, which collects more than 200 deep mutational scanning assays and evaluates both zero-shot and supervised predictors on these datasets.^7^ Both resources primarily assess how well a model predicts or ranks variants in a single pass, rather than how a campaign performs when the model is iteratively retrained to select successive batches of variants for experimental testing.

The second group examines individual design choices within a campaign. Regarding protein representations, Biswas et al. showed that language-model embeddings allow useful predictors to be fitted from a few dozen measurements,^8^ and Hsu et al. found that combining evolutionary density models with assay labels outperformed either source alone.^9^ For initial training-set selection, Wittmann et al. showed using a four-site library that the initial training set largely determines whether a campaign reaches the global optimum, and that zero-shot prefiltering improves the outcome,^3^ while the CLADE approach by Qiu and Wei uses clustering to sample diverse variants.^10^ Regarding acquisition strategies and batch size, Hu et al. compared batch sizes,^11^ and Cheng et al. compared batch sizes and acquisition functions.^12^ Yang et al. compared acquisition functions with the representation fixed.^13^ These studies examine selected campaign design choices. Here we evaluate representation, regression model, acquisition rule, and batch size together.

The third group compares strategies across landscapes. Li et al. evaluated several MLDE strategies on 16 combinatorial landscapes and related their performance to landscape properties, such as navigability.^14^ Together, these studies demonstrate that MLDE strategies can be compared systematically, but an important question remains: if a benchmark identifies the best combination of representation and prediction model, will that combination also perform well on a previously unseen protein? Answering this requires a systematic evaluation of multiple interacting design choices across diverse experimental landscapes, particularly whether their relative performance remains consistent across landscapes.

Here, we introduce MLDEx, a large-scale benchmark comprising 557,143 simulated MLDE campaigns across 16 experimentally characterized protein fitness landscapes. We systematically vary five aspects of campaign design: protein representation, regression model, acquisition rule, initial training-set size, and acquisition batch size. We first quantify the benefit of ML guidance compared with random and zero-shot screening, the latter ranking variants using a pretrained model without experimental fitness measurements. We then examine how acquisition strategy, batch size, representation, and regression model influence campaign performance, and compare the computational cost of leading configurations. Finally, we test whether configuration choices identified from a benchmark of multiple landscapes retain their performance on a previously unseen landscape, and whether alternative selection criteria provide more reliable recommendations.

Across the 16 landscapes, ML-guided campaigns provided a substantial advantage over uniform random screening when the initial training set was small, although this advantage decreased as the initial set increased. Furthermore, greedy acquisition and smaller acquisition batches early in a campaign offered broadly consistent benefits. By contrast, configurations selected using 15 landscapes achieved success rates that were, on average, 13.3 percentage points lower on the held-out landscape than on the landscapes used for selection. These findings provide practical guidance for interpreting MLDE benchmarks and designing protein-engineering campaigns.

## RESULTS

### Overview of the MLDEx benchmark

We established a common framework for comparing ML-guided directed evolution across experimentally measured protein fitness landscapes. The 16 datasets cover four assay classes (binding, activity, stability, and organismal fitness) and comprise 11 combinatorial libraries and five substitution scans, ranging from 2,638 to 536,962 measured variants (**Figure 1a** and **Table 1**). MLDE campaigns were simulated in which a fitness predictor was iteratively trained, used to select additional variants, and updated with their experimental fitness values. Of the 557,143 completed campaigns, 516,068 were retained for the principal analyses after excluding Gaussian-process configurations with uneven coverage across landscapes (**Figure 1b**).

**Figure 1.**
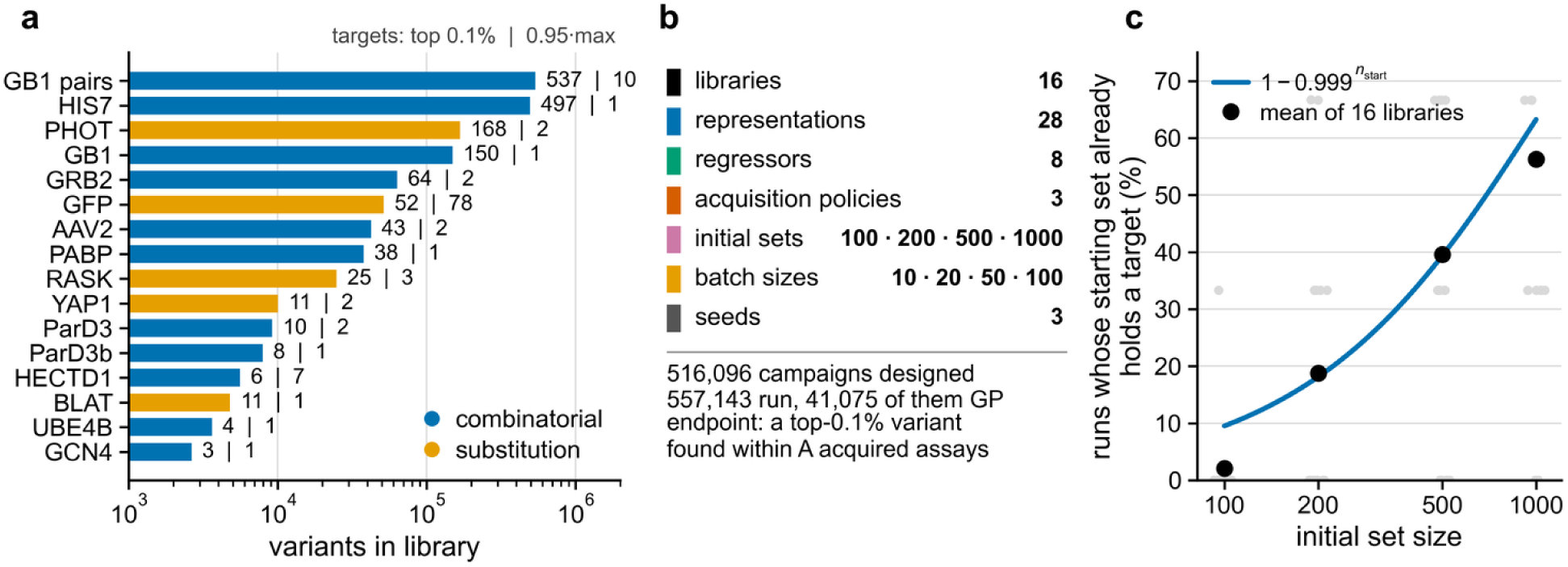
Design of the MLDEx benchmark and the effect of the initial training set. (a) The 16 experimental fitness landscapes, ordered by number of measured variants (**Table 1** and **Table S1**). Blue bars denote combinatorial libraries and orange bars substitution scans. Numbers indicate how many variants meet the top-0.1% success criterion and the separate criterion of fitness at least 95% of the landscape maximum. (b) Combinations of protein representations, regression models, acquisition strategies, initial training-set sizes, batch sizes, and random seeds assessed in the simulations. The analysis excludes the unevenly completed Gaussian-process runs. (c) Percentage of initial training sets that already contain a top-0.1% variant, as a function of starting-set size. Grey points show individual landscapes, black points their mean, and the curve the theoretical expectation for uniform sampling of a landscape with approximately 0.1% target variants. Such campaigns are excluded from comparisons between MLDE configurations but retained in comparisons with screening.

**Table 1.** Experimental fitness landscapes included in MLDEx. Landscapes are grouped by library construction and list the measured property, number of variants, and number of variants reaching the top-0.1% fitness threshold or at least 95% of the maximum fitness. Success at 100 and 400 acquired assays denotes the percentage of eligible campaigns using greedy acquisition that identified a top-0.1% variant. We included all variants tied at the highest-fitness 0.1% boundary, so BLAT has 11 rather than five successful variants. The Source column cites the original dataset studies. **Table S1** gives the ProteinGym accessions.

| Landscape | Assay | # Variants | # Target Variants |  | Success (%) |  | Source |
|---|---|---|---|---|---|---|---|
|  |  |  | top 0.1% | 95% max | 100 assays | 400 assays |  |
| *Combinatorial* |  |  |  |  |  |  |  |
| GCN4 | binding | 2,638 | 3 | 1 | 15 | 40 | Staller *et al.*<sup>15</sup> |
| UBE4B | stability | 3,622 | 4 | 1 | 24 | 51 | Tsuboyama *et al.*<sup>16</sup> |
| HECTD1 | stability | 5,586 | 6 | 7 | 63 | 86 | Tsuboyama *et al.*<sup>16</sup> |
| ParD3b | organismal fitness | 7,922 | 8 | 1 | 73 | 97 | Ding *et al.*<sup>17</sup> |
| ParD3 | organismal fitness | 9,192 | 10 | 2 | 85 | 95 | Aakre *et al.*<sup>18</sup> |
| PABP | organismal fitness | 37,708 | 38 | 1 | 37 | 72 | Melamed *et al.*<sup>19</sup> |
| AAV2 | organismal fitness | 42,328 | 43 | 2 | 32 | 61 | Sinai *et al.*<sup>20</sup> |
| GRB2 | organismal fitness | 63,366 | 64 | 2 | 40 | 75 | Faure *et al.*<sup>21</sup> |
| GB1 | binding | 149,360 | 150 | 1 | 75 | 93 | Wu *et al.*<sup>22</sup> |
| HIS7 | organismal fitness | 496,137 | 497 | 1 | 18 | 44 | Pokusaeva *et al.*<sup>23</sup> |
| GB1 pairs | binding | 536,962 | 537 | 10 | 56 | 78 | Olson *et al.*<sup>24</sup> |
| *Substitution scan* |  |  |  |  |  |  |  |
| BLAT | organismal fitness | 4,783 | 11 | 1 | 53 | 76 | Firnberg *et al.*<sup>25</sup> |
| YAP1 | binding | 10,075 | 11 | 2 | 68 | 88 | Araya *et al.*<sup>26</sup> |
| RASK | binding | 24,873 | 25 | 3 | 40 | 68 | Weng *et al.*<sup>27</sup> |
| GFP | activity | 51,714 | 52 | 78 | 28 | 58 | Sarkisyan *et al.*<sup>28</sup> |
| PHOT | activity | 167,529 | 168 | 2 | 93 | 98 | Chen *et al.*<sup>29</sup> |

Campaign success was defined as identifying at least one variant among the top 0.1% of the corresponding fitness landscape. This percentile-based criterion provides a consistent measure across landscapes, whereas a threshold based on a fixed fraction of the maximum fitness would yield highly variable numbers of successful variants. For example, requiring at least 95% of the maximum fitness would identify only one or two qualifying variants in 12 of the 16 landscapes (**Table 1**). Campaign success was therefore evaluated after fixed numbers of newly acquired variants (e.g., 100 or 400), rather than only at campaign termination. This ensured that campaigns with different batch sizes were compared at the same number of newly acquired experimental assays.

An additional consideration was that successful variants could already be present in the initial training set before ML guidance began. Although variants at or above the separate 95%-of-maximum stopping threshold were excluded from initial sampling, not all variants within the top 0.1% were excluded. Among campaigns initialized with 1,000 variants, a top-0.1% variant was already present in 56.2% of cases, compared with a theoretical probability of approximately 63.2% under fully uniform sampling (**Figure 1c**). Because these campaigns succeeded before model training, no information about the effectiveness of model-guided selection could be obtained from them. They were therefore excluded from comparisons between MLDE configurations, leaving 365,563 campaigns for these analyses. However, they were retained in comparisons with random and zero-shot screening, where initial measurements were included in the total assay budget for all approaches.

### ML guidance improves screening efficiency, especially with small initial training sets

We next asked whether iterative ML-guided selection identifies highly functional variants more efficiently than screening without guidance. For this analysis, MLDE campaigns with greedy acquisition were examined. Rather than focusing on a particular prediction model, all eligible combinations of the 28 protein representations and eight regression models were considered across the different acquisition batch sizes. Success rates were averaged across these configurations to assess the overall benefit of ML guidance. The results were compared with uniform random screening and direct screening of variants ranked by a pretrained structure-conditioned model. Importantly, comparisons were performed at matched total assay budgets, including the variants used to train the initial ML model. The analytical random-screening baseline reached 50% success at approximately 621 assays (**Figure 2a**) (“Screening controls and additional experiments”).

**Figure 2.**
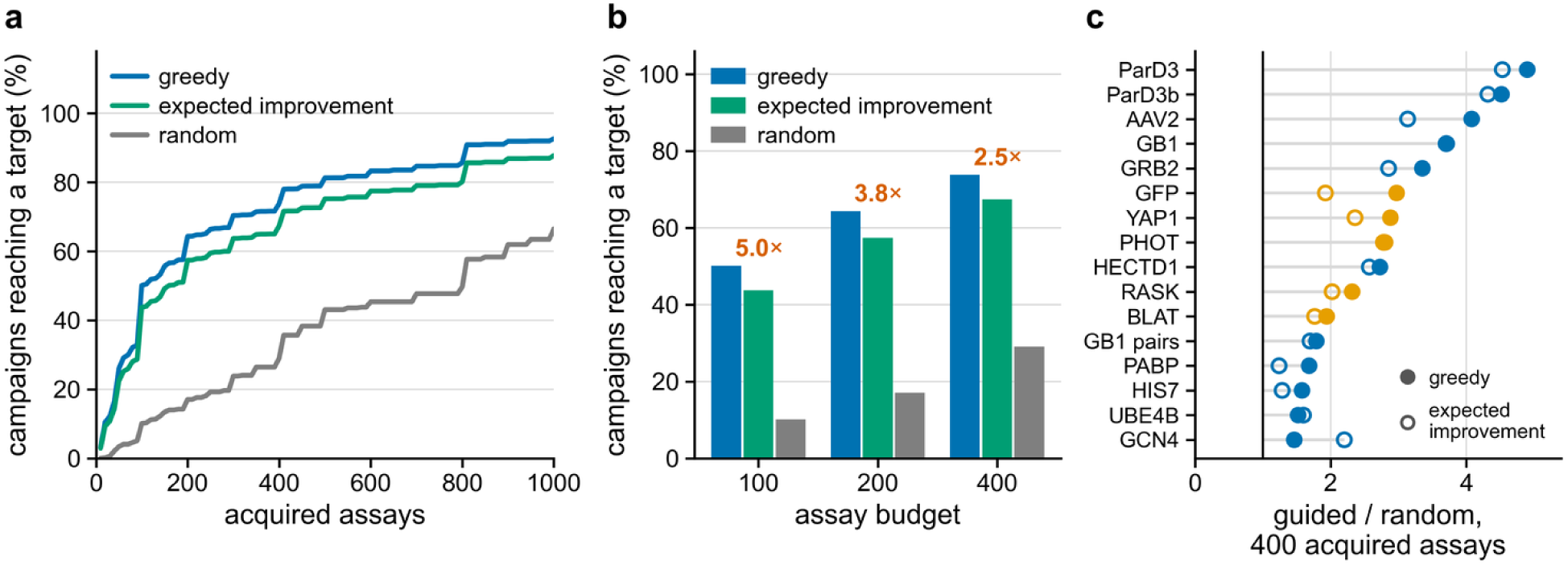
Comparison of ML-guided variant selection with random screening. (a) Landscape-balanced percentage of campaigns finding a top-0.1% variant as additional variants are acquired using greedy selection, expected improvement, or uniform random acquisition. This panel counts only acquisitions after the initial training set: greedy acquisition reaches 50% success after 100 newly acquired assays, expected improvement after 160, simulated random acquisition after 810 assays, and unrestricted analytical random screening after approximately 621 assays (“Screening controls and additional experiments”). (b) Guided and random-acquisition success at 100, 200, and 400 newly acquired assays. Numbers above the bars indicate the ratio of their success rates. (c) Improvement from greedy acquisition over random acquisition in each landscape after 400 newly acquired assays. When the initial training measurements are included, the advantage is smaller (Figure 3).

A clear advantage of ML guidance over uniform random screening was observed when initial training sets were small. With 100 starting variants and 100 subsequently acquired variants (i.e. a total of 200 experimental measurements) a top-0.1% variant was identified in 44.6% of ML-guided campaigns, compared with 19.9% for random screening. This corresponds to a 2.24-fold higher success probability. Across the smaller initial training-set sizes, a given success level was achieved with approximately 2.2–2.8 times fewer total assays using ML guidance than random screening. However, this advantage diminished as the initial training-set size increased. With 1,000 initial variants, the success ratio declined to 1.12 at 1,100 total assays, across the 11 landscapes with eligible runs under this condition (**Figures 2** and **3a**).

**Figure 3.**
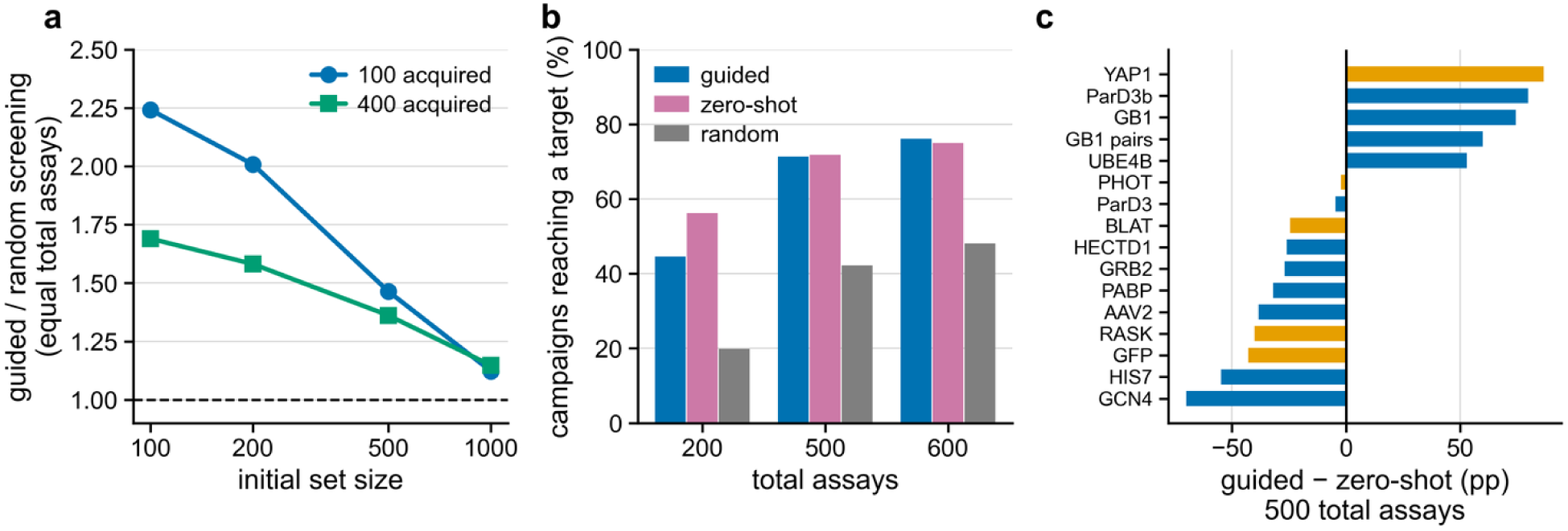
Effect of initial training-set size on the benefit of ML guidance. (a) Ratio of ML-guided to uniform random-screening success rates at matched total assay budgets, shown for different initial training-set sizes after 100 or 400 additional assays. (b) Success rates of ML-guided campaigns, ESM-IF1-based zero-shot screening,^30^ and uniform random screening at total assay budgets of 200 (100 initial + 100 acquired), 500 (100 + 400), and 600 (200 + 400) assays. (c) Difference in success rates between ML-guided campaigns and ESM-IF1-based zero-shot screening across individual landscapes at a total budget of 500 assays. Positive values indicate an advantage for ML guidance. Blue and orange bars represent combinatorial libraries and substitution scans, respectively.

This decline can be explained by the increasing experimental effort required to generate the initial training set. Measuring 1,000 variants already represents a substantial random-screening experiment, with a high probability of identifying a successful variant before model training begins. Consequently, a smaller additional benefit can be obtained from subsequent ML-guided selection when all experimental measurements are considered. In contrast, when only newly acquired variants were counted, a considerably greater advantage of ML guidance was observed (**Figure 2b**). After 400 acquired assays, greedy acquisition outperformed random acquisition on all 16 landscapes, with a median 2.8-fold advantage that ranged from 1.5-to 4.9-fold across landscapes (**Figure 2c**). Both comparisons are informative, but only the total-assay comparison accounts for the full experimental effort required.

A stronger alternative to uniform random screening is zero-shot screening, in which variants are ranked by a pretrained protein model without using experimentally measured fitness values. Here, ESM-IF1, a structure-conditioned model, was used to rank variants according to the average log-probability assigned to their mutated residues. These scores showed a mean Spearman correlation of ρ = 0.37 with experimental fitness across the 16 landscapes (“Screening controls and additional experiments”; **Result S1**). Unlike iterative MLDE, zero-shot screening does not require a campaign-specific prediction model to be trained.

At 200 total assays, a successful variant was identified in 56.2% of cases using zero-shot screening, compared with 44.6% for ML-guided campaigns. Higher success rates with ML guidance were observed on seven of the 16 landscapes. However, the difference diminished at larger assay budgets. At 500 assays, success rates of 71.3% and 71.9% were obtained for zero-shot and ML-guided screening, respectively, while at 600 assays the corresponding rates were 76.1% and 75.0%. Higher success rates with ML guidance were observed on five of the 16 landscapes at each of these larger budgets (**Figure 3b,c**). Thus, under the tested conditions, comparable performance was achieved by zero-shot screening and iterative ML-guided selection at matched total assay budgets.

These results should not be interpreted as evidence that MLDE is generally inferior to zero-shot screening. In the present benchmark, all ML-guided campaigns were initialized with uniformly sampled variants, whereas zero-shot screening allowed promising variants to be prioritized without requiring initial experimental measurements. The importance of initial training-set selection is therefore highlighted by this comparison. Further improvements in MLDE performance may be achieved by combining zero-shot prioritization with MLDE initialization, although this strategy was not evaluated here.

### Effects of acquisition strategy, batch size, representation, and regression model

We next examined the individual design choices that define an MLDE campaign: the strategy used to select variants, the number of variants tested between model updates, the representation used to encode protein sequences, and the regression model used to predict fitness. The 28 representations included one-hot and BLOSUM encodings, physicochemical descriptors, sequence- and structure-informed protein models, and representations restricted to mutated positions or expressed as differences from the wild-type sequence (**Table S2**). The eight evaluated regression models comprised linear regression, ridge regression, elastic net, SVR, random forest, LightGBM, XGBoost, and MLP (**Table S4**). The effects of these design choices were compared across the 16 landscapes after 100 and 400 newly acquired assays, with an initial focus on identifying strategies that performed consistently across different proteins. **Figure S3** shows the design-factor effects by landscape.

Greedy acquisition was more effective than expected improvement. After 400 acquired assays, greedy acquisition increased the success rate by 6.3 percentage points (pp) (95% CI, 5.5–7.2; exact sign-flip p = 0.0211; wild-cluster p < 0.0001). A similar difference of 6.2 pp was observed after 100 acquired assays. Greedy acquisition outperformed expected improvement on 14 of the 16 landscapes at both assay counts, and successful variants were identified an average of 21.2 assays earlier (95% CI, 18.0–24.6) (**Figure 4d**; **Table 2**). The two landscapes favoring expected improvement were the smallest in the benchmark.

**Figure 4.**
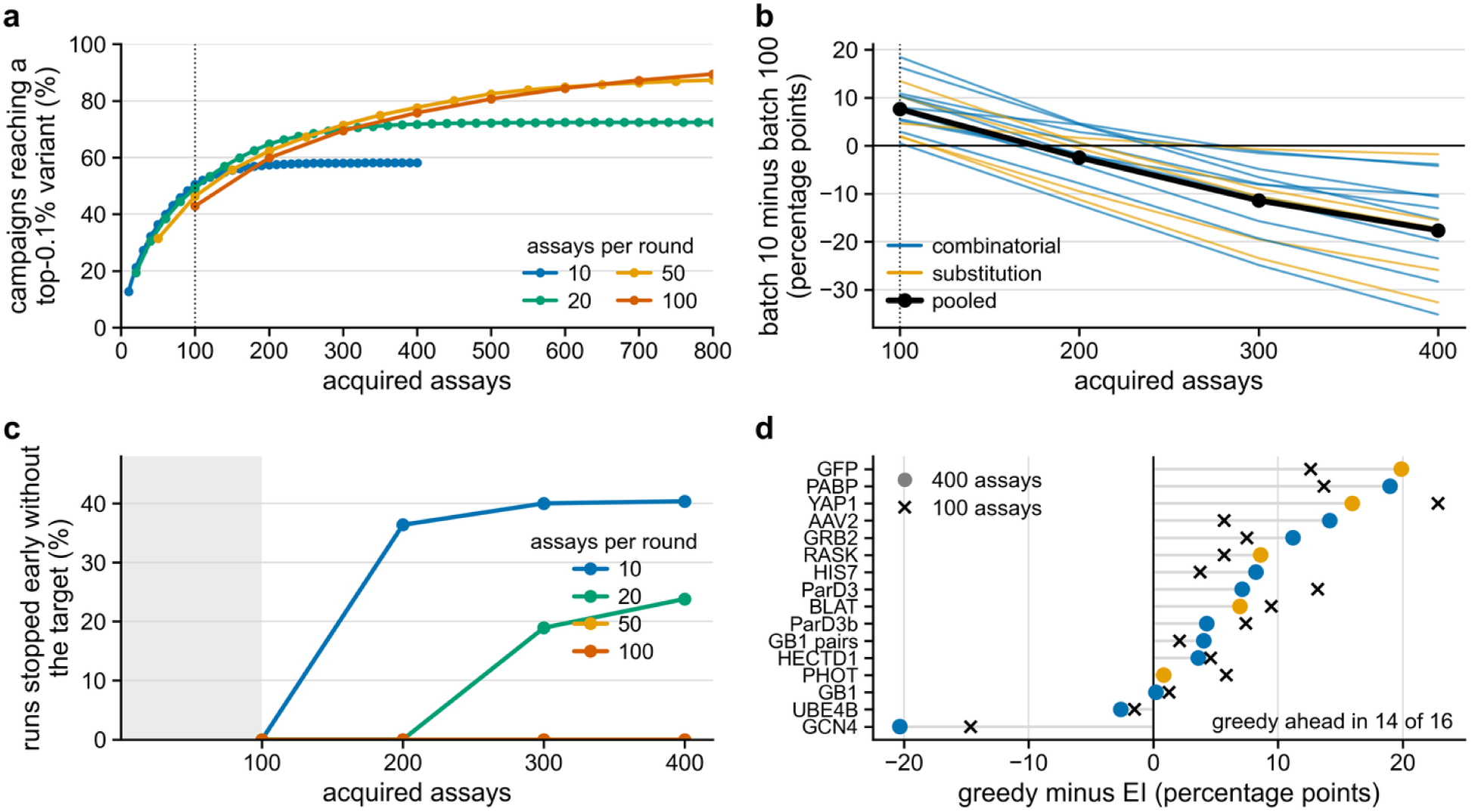
Effects of acquisition strategy and batch size on the identification of high-fitness variants. (a) Landscape-balanced success rates as a function of the number of newly acquired assays for batch sizes of 10, 20, 50, and 100. The dotted line indicates 100 acquisitions, before the early-stopping rule can influence the results. (b) Difference in success rates between batch sizes of 10 and 100 across landscapes. Positive values indicate an advantage for smaller batches. Blue and orange lines represent combinatorial libraries and substitution scans, respectively, while the black line shows the landscape-balanced mean. (c) Percentage of runs stopped early without finding a target, by the number of acquired assays and batch size. Ten rounds correspond to 100 acquisitions for batches of 10 but 1,000 acquisitions for batches of 100, allowing campaigns with smaller batches to terminate earlier. (d) Difference in success rates between greedy and expected-improvement acquisition after 100 and 400 newly acquired assays across individual landscapes.

**Table 2.**
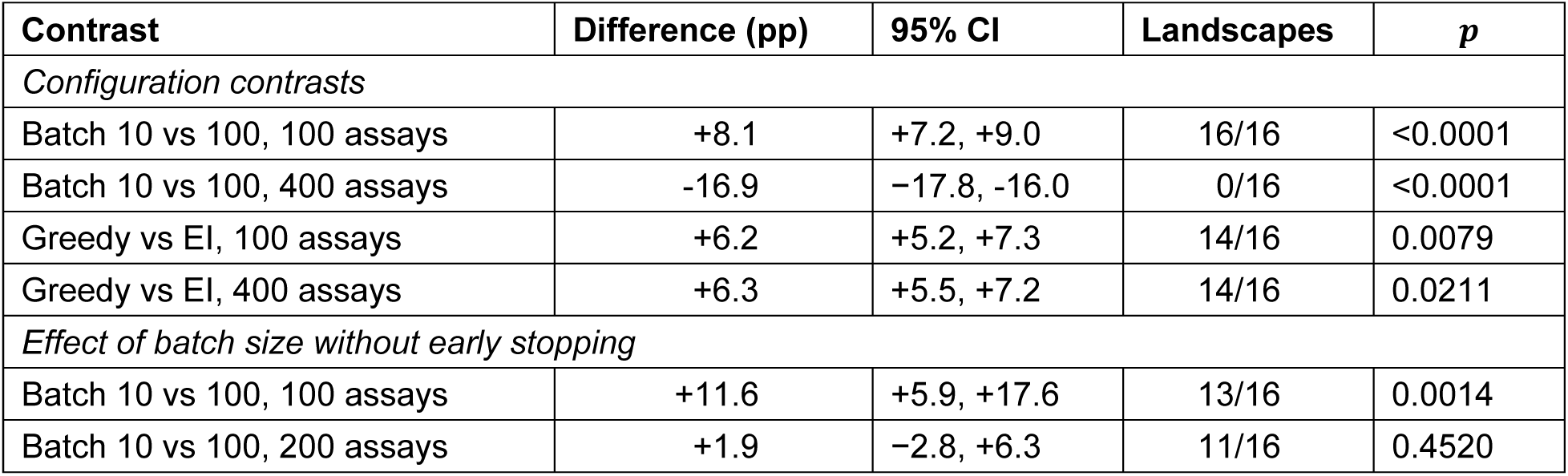
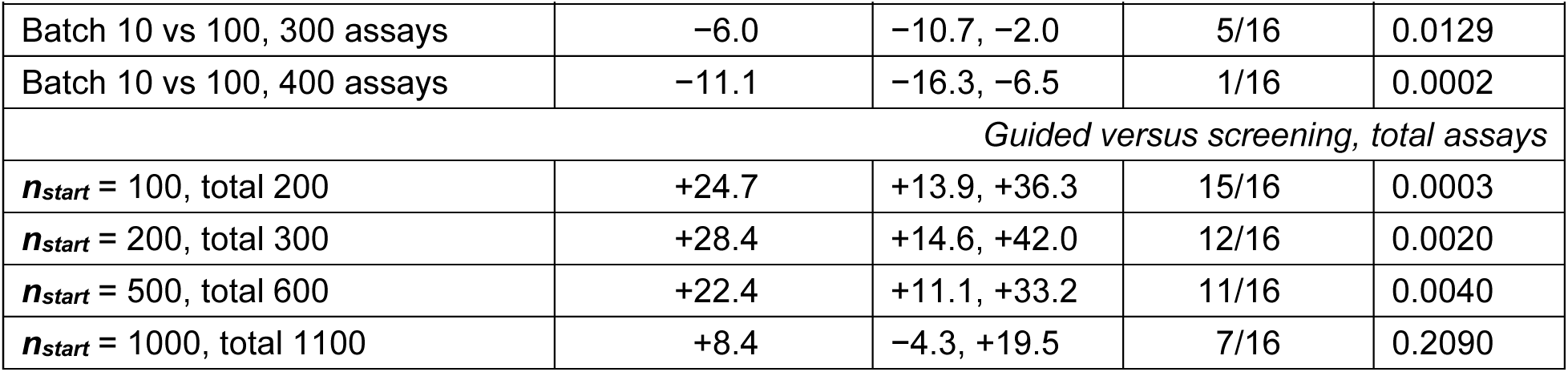
Key comparisons between MLDE strategies across protein fitness landscapes. Values represent landscape-balanced differences in the probability of identifying a top-0.1% variant, with 95% confidence intervals. Comparisons between MLDE configurations exclude campaigns that were already successful at initialization, whereas comparisons with random screening include the initial training set in the total assay budget. Confidence intervals were calculated by resampling scenarios for acquisition strategy and batch-size comparisons, and by resampling landscapes for the experiment without early stopping. The number of landscapes favoring each strategy and the corresponding exact sign-flip p values are also reported. A separate landscape-level bootstrap yielded consistent estimates for the four main protocol comparisons.

| Contrast | Difference (pp) | 95% CI | Landscapes | *p* |
|---|---|---|---|---|
| *Configuration contrasts* |  |  |  |  |
| Batch 10 vs 100, 100 assays | +8.1 | +7.2, +9.0 | 16/16 | <0.0001 |
| Batch 10 vs 100, 400 assays | -16.9 | −17.8, -16.0 | 0/16 | <0.0001 |
| Greedy vs EI, 100 assays | +6.2 | +5.2, +7.3 | 14/16 | 0.0079 |
| Greedy vs EI, 400 assays | +6.3 | +5.5, +7.2 | 14/16 | 0.0211 |
| *Effect of batch size without early stopping* |  |  |  |  |
| Batch 10 vs 100, 100 assays | +11.6 | +5.9, +17.6 | 13/16 | 0.0014 |
| Batch 10 vs 100, 200 assays | +1.9 | −2.8, +6.3 | 11/16 | 0.4520 |
| Batch 10 vs 100, 300 assays | −6.0 | −10.7, −2.0 | 5/16 | 0.0129 |
| Batch 10 vs 100, 400 assays | −11.1 | −16.3, −6.5 | 1/16 | 0.0002 |
| *Guided versus screening, total assays* |  |  |  |  |
| ***n***<sub>***start***</sub> = 100, total 200 | +24.7 | +13.9, +36.3 | 15/16 | 0.0003 |
| ***n***<sub>***start***</sub> = 200, total 300 | +28.4 | +14.6, +42.0 | 12/16 | 0.0020 |
| ***n***<sub>***start***</sub> = 500, total 600 | +22.4 | +11.1, +33.2 | 11/16 | 0.0040 |
| ***n***<sub>***start***</sub> = 1000, total 1100 | +8.4 | −4.3, +19.5 | 7/16 | 0.2090 |

The number of variants tested between model updates also influenced campaign performance. During the first 100 acquisitions, batches of 10 variants achieved an 8.1-percentage-point higher success rate than batches of 100 (95% CI, 7.2–9.0), with smaller batches performing better on all 16 landscapes (**Figure 4a,b**; **Table 2**). After 400 acquired assays, however, the difference was reversed (−16.9 pp), favoring larger batches. Because campaigns could be terminated after ten consecutive rounds without improvement, this late-stage difference may partly reflect the early-stopping rule rather than batch size alone (**Figure 4c**).

To distinguish the effects of batch size from early stopping, an additional set of 4,608 campaigns was simulated in which the early-stopping threshold was increased to 40 consecutive rounds without improvement, preventing termination due to this criterion before 400 variants had been acquired. The experiment included all 16 landscapes, an initial training set of 100 variants, four representations, three regression models, two acquisition rules, and three random seeds. Each campaign was designed to acquire 400 additional variants (“Screening controls and additional experiments”). The reversal in batch-size performance persisted under these conditions (**Figure 5a**). After 100 acquired assays, batches of 10 showed an 11.6 pp advantage over batches of 100 (95% CI, 5.9–17.6), with smaller batches performing better on 13 of 16 landscapes. At 200 assays, the difference was small and not statistically significant (1.9 pp; 95% CI, −2.8 to 6.3).

**Figure 5.**
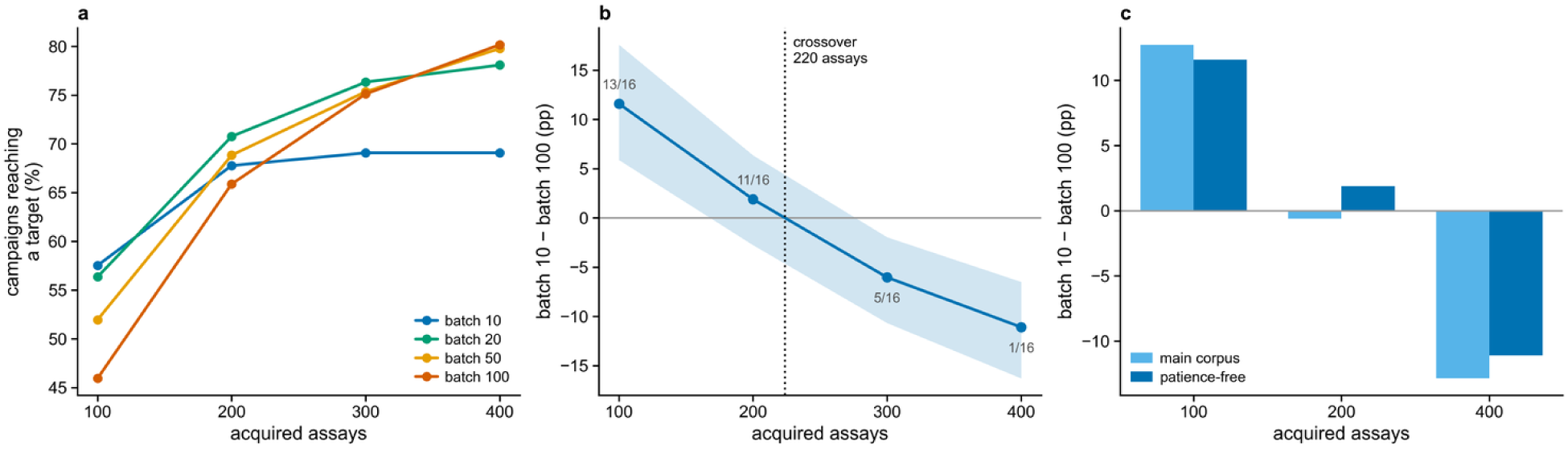
The benefit of small acquisition batches reverses even when early stopping is disabled. All campaigns in this additional experiment were designed to acquire 400 variants. (a) Landscape-balanced success as a function of acquired assays for each batch size. (b) Difference in success between batches of 10 and 100, with confidence intervals obtained by resampling landscapes; positive values favor batches of 10. Numbers indicate how many landscapes favor the smaller batch. The dotted line marks the crossover near 220 acquired assays. (c) Comparison of the batch-size effect in the main benchmark and in the additional experiment, using scenarios available in both analyses.

After 400 assays, batches of 100 instead achieved an 11.1 pp higher success rate (95% CI, 6.5–16.3), with smaller batches performing better on only one landscape. In matched scenarios, the differences between batches of 10 and 100 were 12.7 pp versus 11.6 pp at 100 assays in the original and additional experiments, respectively, and −12.9 pp versus −11.1 pp at 400 assays (**Figure 5c**; **Table 2**). Interpolation indicated a crossover at approximately 224 newly acquired variants (**Figure 5b**), compared with approximately 176 acquisitions in the original experiment (**Result S2; Figures S4** and **S5**). These results indicate that frequent model updates are most beneficial early in a campaign, whereas larger batches may be preferable when more variants are tested.

The choice of regression model was another important determinant of success, although the relative influence of individual design choices varied across landscapes (**Figure S3**). The MLP regressor performed worst among the eight evaluated models on 14 of the 16 landscapes after both 100 and 400 acquired assays. At 400 assays, a landscape-balanced success rate of 53.6% was obtained with MLP, compared with 81.5% for XGBoost (**Figure 6**; **Figure S9**). Thus, the poor performance of MLP was remarkably consistent across landscapes, whereas no individual representation was the worst-performing option on more than six landscapes.

**Figure 6.**
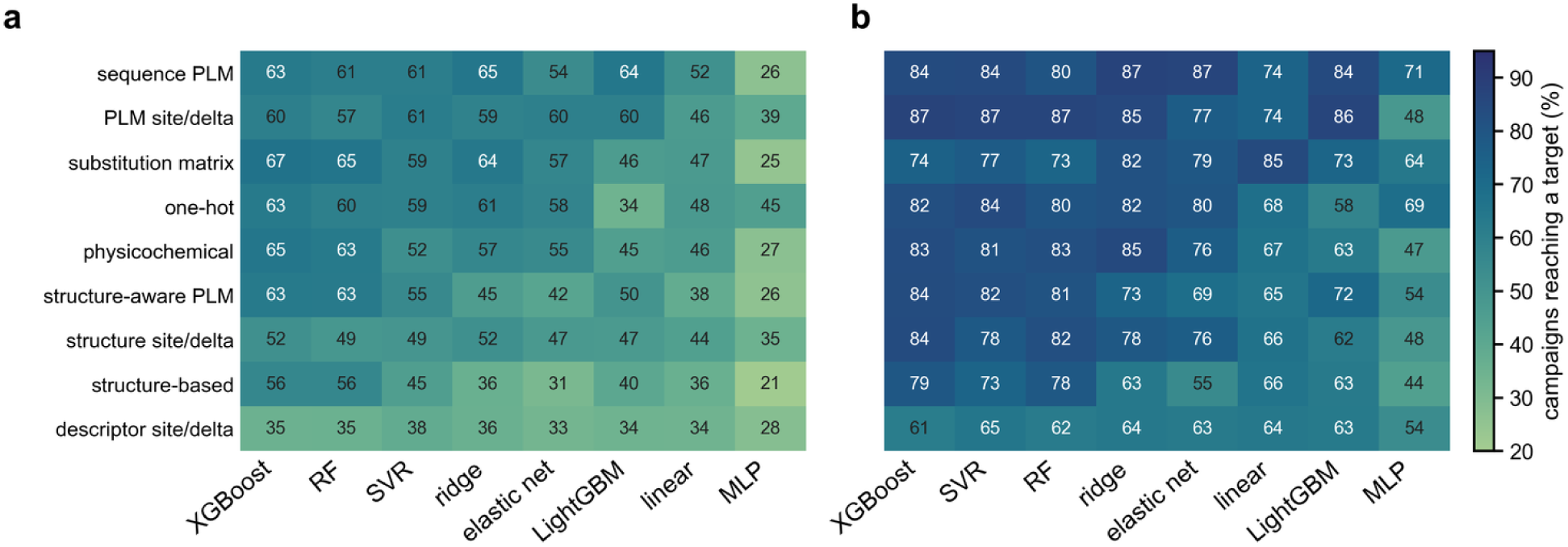
Success rates for combinations of representation families and regression models. Heatmaps show the landscape-balanced proportion of campaigns identifying a top-0.1% variant after (a) 100 and (b) 400 acquired assays using greedy acquisition. Columns represent the eight regression models, while rows group the 28 protein representations by family. Sequence PLM includes sequence-based protein language models; structure-aware PLM includes models such as ProstT5, SaProt, ProSST, and VenusREM; and structure-based representations include ESM-IF1, ProteinMPNN probabilities, and SLAE. “Site/delta” denotes representations restricted to mutated positions or calculated as differences between variant and wild-type features. The individual representations are listed in **Table S2**. Rows and columns are ordered by mean performance after 100 acquisitions. The relative success rates depend strongly on the combination of representation and regression model.

Campaign performance also depended on the combination of representation and regression model (**Figure 6a,b**; **Figures S6–S9**). No individual representation–model combination ranked among the bottom 10% on more than ten landscapes at 400 acquired assays. Across all 224 combinations, 28.8% of the variation in success was attributed to the representation (95% CI, 21.6–38.3%), 33.2% to the regression model (22.8–42.6%), and 38.0% to their interaction (33.2–44.0%). Similar contributions were observed after 100 acquisitions. These findings indicate that the performance of representations and regression models cannot be fully assessed independently, as their combination substantially influences campaign success.

The benefit of protein language model representations over simpler encodings, such as one-hot, BLOSUM, and physicochemical descriptors, was also examined. Averaged across regression models, protein language model representations increased success by 5.2 pp after 400 acquired assays (95% CI, 4.3–5.9), but by only 0.8 pp after 100 assays (95% CI, −0.1 to 1.9; **Figure 7a**). However, their benefit varied considerably between regression models. With LightGBM, protein language model representations increased success by 20.3 pp and outperformed simpler encodings on 13 of 16 landscapes (exact sign-flip p = 0.0007). In contrast, an improvement of only 0.6 pp was observed with ridge regression (**Figure 7b**). At 100 assays, simpler encodings performed better with ridge regression (−3.2 pp), elastic net (−3.6 pp), random forest (−3.7 pp), and XGBoost (−3.6 pp), where negative values indicate an advantage for simpler encodings over PLMs.

**Figure 7.**
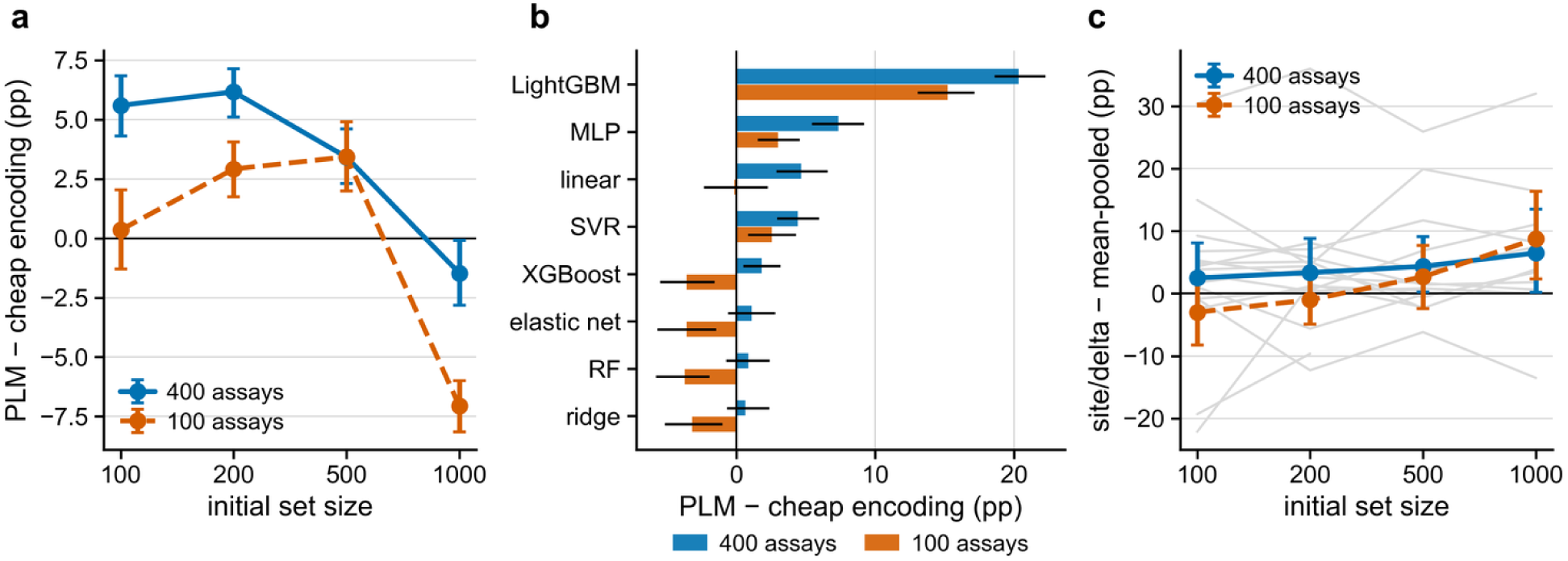
The benefit of protein language model representations depends on the regression model and the amount of training data. Landscape-balanced differences in success rates between protein language model representations and simpler encodings (one-hot, substitution matrices, and physicochemical descriptors). Positive values indicate an advantage for protein language model representations. (a) Differences by initial training-set size. (b) Differences across regression models. (c) Comparison of full-sequence mean-pooled embeddings with representations restricted to mutated positions. All comparisons were paired within the same scenario and regression model.

The initial training-set size also influenced the benefit of protein language model representations. Their average advantage decreased from 5.6 pp with 100 initial variants to −1.5 pp with 1,000 variants (**Figure 7a**). The heatmaps in **Figure 6** and **Figures S6–S9** further illustrate the dependence of representation performance on the regression model. In these analyses, “sequence PLM” refers to sequence-based protein language model representations (e.g., ESM-2, ProtT5, Ankh, and ESM-C), whereas “structure-aware PLM” includes models incorporating structural information (e.g., ProstT5, SaProt, ProSST, and VenusREM). “Structure-based” refers to representations derived from ESM-IF1, ProteinMPNN, or SLAE. Labels ending in “site/delta” denote representations restricted to mutated positions (“site”; **Figure 7c**) or calculated as differences between variant and wild-type features (“delta”). This convention was applied to PLM, structure-derived, and descriptor representations. All individual representations, including the eight site/delta encodings, are listed in **Table S2**. Together, these findings help reconcile previous reports of substantial improvements in MLDE performance using protein language model embeddings^8^ with evidence from fitness-prediction benchmarks that simpler encodings can be competitive.^6^ The study by Hsu et al.^9^ addresses the related problem of learning fitness models from evolutionary and assay-labeled data. Both outcomes were observed in the present benchmark, depending on the regression model and the amount of available training data.

### Computational cost distinguishes similarly successful configurations

We next compared the performance and computational requirements of all 224 representation–regression-model combinations. Computational cost is an important consideration in practical protein engineering, as certain configurations may require substantially more computing time without achieving higher experimental success rates. At 400 acquired assays, 26 combinations could not be statistically distinguished from the highest-performing combination using a paired bootstrap, although their model-fitting times differed by a factor of 1,043 (**Figure 8b**). The highest observed success rate (89.7%) was achieved with ProstT5 and random forest, requiring 2.857 CPU-hours per campaign. In comparison, a mutation-site-restricted SLAE representation combined with linear regression (site:SLAE + linear) achieved 87.5% success with only 0.005 CPU-hours, without a statistically significant difference from the leading combination. At 100 acquired assays, only three combinations were statistically indistinguishable from the leading Ankh-base + ridge model, which achieved a success rate of 73.5% (**Figure 8a**).

**Figure 8.**
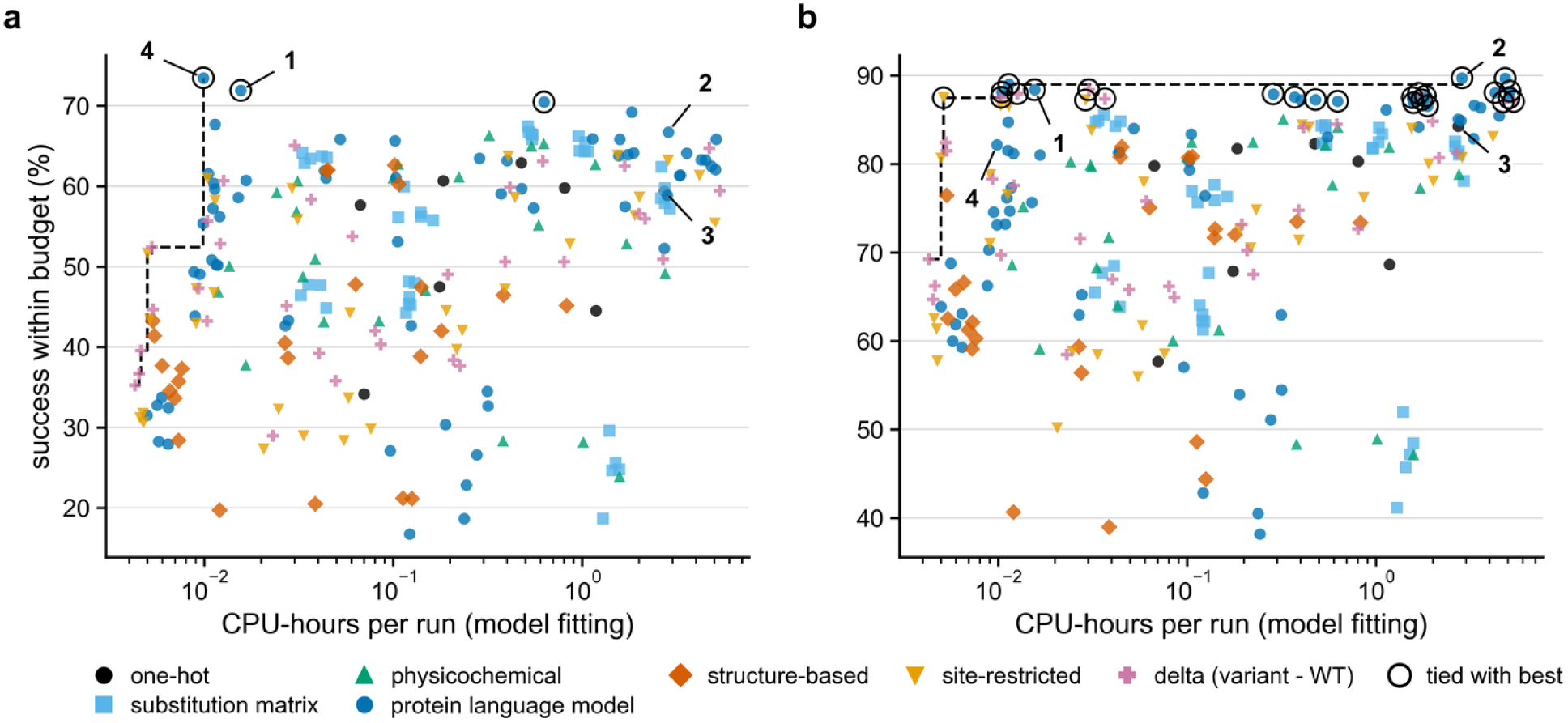
Trade-off between success rate and regression-model fitting time. Each point represents one of the 224 representation–regression-model combinations under greedy acquisition. The horizontal axis shows mean CPU-hours spent fitting models per campaign and the vertical axis the landscape-balanced success rate after (a) 100 or (b) 400 newly acquired assays. Open circles mark combinations whose success was not statistically distinguishable from the best observed combination (three at 100 assays and 26 at 400 assays). The dashed line shows the best observed success achievable at each computational cost. Numbered examples are 1, SaProt + ridge; 2, ProstT5 + random forest; 3, one-hot + SVR; and 4, Ankh-base + ridge. The inexpensive site:SLAE + linear combination achieved 87.5% success at 0.005 CPU-hours in panel b. Plot symbols group representations as one-hot, substitution matrices, physicochemical descriptors, protein language models, structure-based models, mutation-site-restricted features, or variant-minus-wild-type features. Representation names are listed in **Table S2** and regression models in **Table S4**.

Computational cost therefore provides an additional criterion for selecting among configurations with comparable success rates. **Figure 9** shows, for each initial training-set size and assay count, the combination with the highest observed success rate and the best-performing combination requiring no more than 0.05 CPU-hours of model fitting per campaign. These comparisons account for model-fitting time but exclude the one-time computational cost of generating sequence embeddings and other representations. The results should therefore be interpreted as descriptive findings of the present benchmark rather than validated recommendations for previously uncharacterized proteins. Furthermore, when many combinations are evaluated, the highest observed success rate may overestimate the performance expected on an independent landscape.^31^

**Figure 9.**
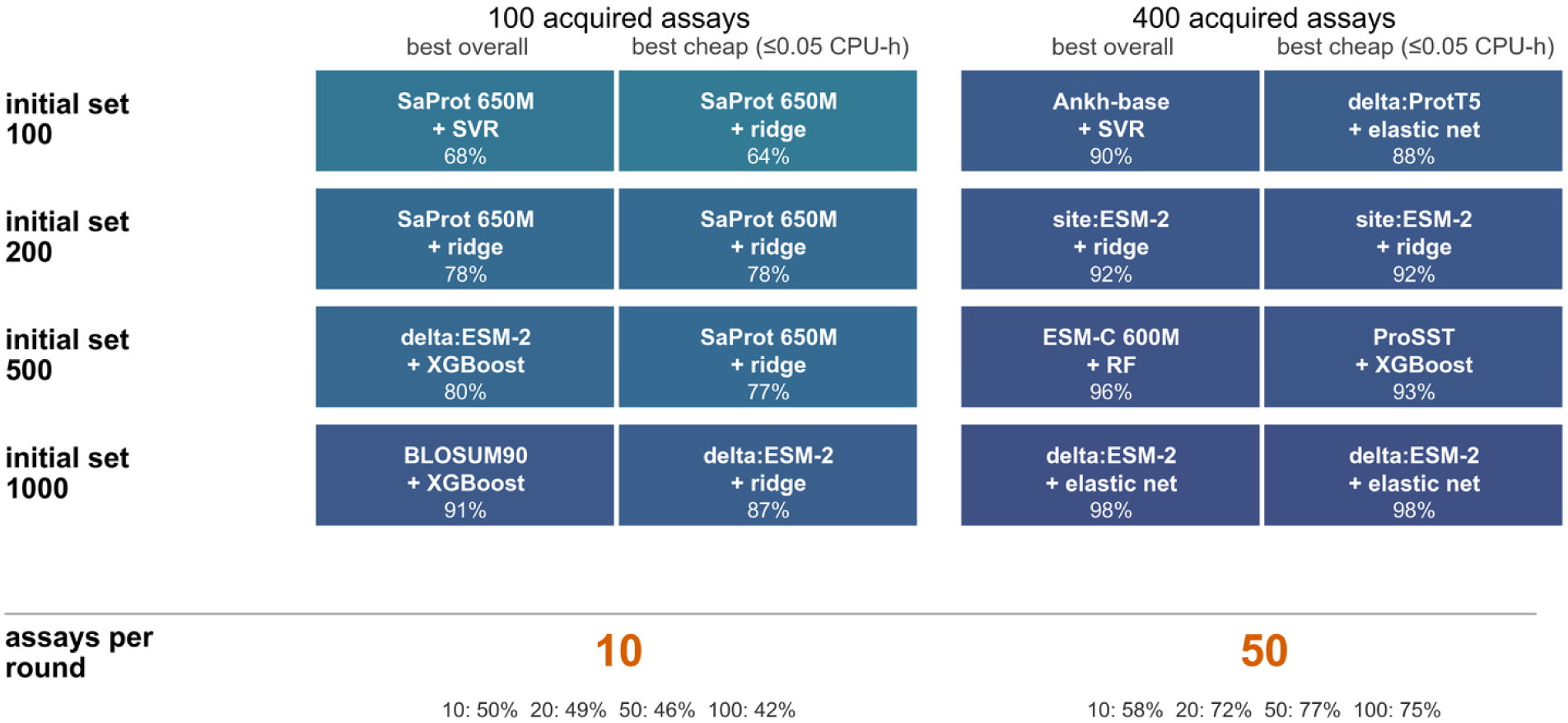
Highest observed success and lower-cost alternatives at different experimental assay counts. For each initial training-set size and number of acquired assays, the summary shows the representation–regression-model combination with the highest landscape-balanced success rate under greedy acquisition (“best overall”) and the best-performing combination requiring no more than 0.05 CPU-hours of model fitting per campaign (“best cheap”). The lower section compares the four acquisition batch sizes at matched assay counts. Because these combinations were selected from a large number of tested configurations, their observed success rates may overestimate performance on previously unseen landscapes. Several configurations achieved statistically indistinguishable success rates (Figure 8), and the rankings of leading representation–model combinations did not reliably transfer to unseen proteins (see next section below).

The best-performing representation–model combination also depended on acquisition batch size. When the 224 combinations were ranked separately for batch sizes of 10, 20, 50, and 100 at 400 acquired assays, a different leading combination was identified for each batch size. Across the four batch sizes, 17 distinct combinations occupied the 20 positions in the respective top-five rankings (**Figure 10**). The highest success rate increased from 84.3% for batches of 10 to 97.2% for batches of 50. However, the mean number of assays required to identify the first successful variant among successful campaigns also increased, from 50 to 161. At 100 acquired assays, the rankings were somewhat more consistent: ten distinct combinations occupied the 20 top-five positions, and Ankh-base + ridge ranked first for three of the four batch sizes (**Figure S10**). Nevertheless, substantial differences between proteins remained, with even the leading combination for batches of 10 achieving success rates below 50% on two landscapes.

**Figure 10.**
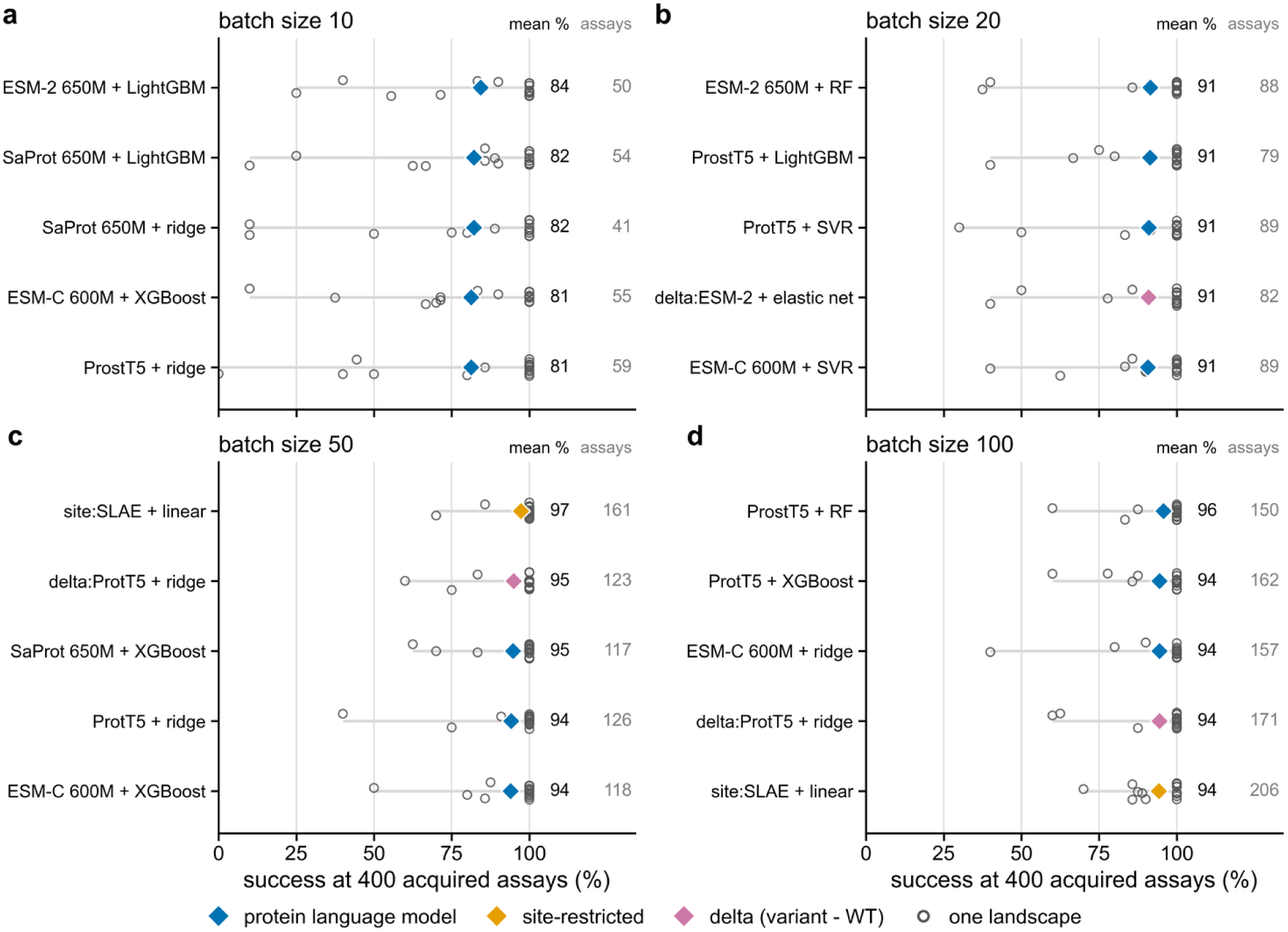
Leading representation–regression-model pairs depend on acquisition batch size. (a–d) Five combinations with the highest landscape-balanced success at 400 acquired assays, ranked separately for batch sizes of 10, 20, 50, and 100 and averaged across the four initial training-set sizes. Each grey range shows success across the 16 landscapes, individual circles show landscape-specific values, and the diamond and adjacent number show the landscape-balanced mean. The right-hand column gives the mean number of acquired assays before the first successful variant was found, among successful campaigns. All 224 combinations were eligible, including the ESM-C representations. Colors indicate the representation families listed in **Table S2**. Corresponding results after 100 acquired assays are shown in **Figure S10**.

### Can a model configuration selected on existing landscapes work on a new protein?

We next evaluated how reliably representation–regression-model combinations selected from an existing benchmark perform on a previously unseen protein fitness landscape. For each of the 16 landscapes, the combination with the highest average campaign success across the remaining 15 landscapes was selected and evaluated on the held-out landscape. This leave-one-landscape-out procedure considered all 224 combinations of 28 representations (**Table S2**) and eight regression models (**Table S4**), using greedy acquisition and matched numbers of newly acquired assays. The performance of their combinations across landscapes is shown in **Figures S6** and **S7** after 100 acquired assays and in **Figure S9** after 400 acquired assays.

Within individual landscapes, configuration rankings were reasonably consistent across replicate simulations. Independent subsets of random seeds yielded a split-half Spearman correlation of ρ = 0.70 after 400 acquired assays. In contrast, substantially lower agreement was observed between landscapes. The mean correlations between landscape-specific rankings were ρ = 0.36 for representations and ρ = 0.43 for regression models, and negatively correlated representation rankings were observed for 15% of landscape pairs (**Figure 11b**). This limited agreement reduced the ability to identify a generally successful configuration. After 400 acquired assays, combinations selected using the other 15 landscapes achieved a median rank of only 93 out of 224 on the held-out landscape and did not rank among the top 10% in any of the 16 held-out evaluations (**Figure 12a**). Their mean success rate on the held-out landscapes was 75.9%, compared with 79.1% for the median combination among the 224 tested configurations (**Figure 11d**). At 100 acquired assays, transfer performance was somewhat better but remained limited, with a median held-out rank of 64 and a top-10% ranking in five of the 16 evaluations (**Figure 11c**).

**Figure 11.**
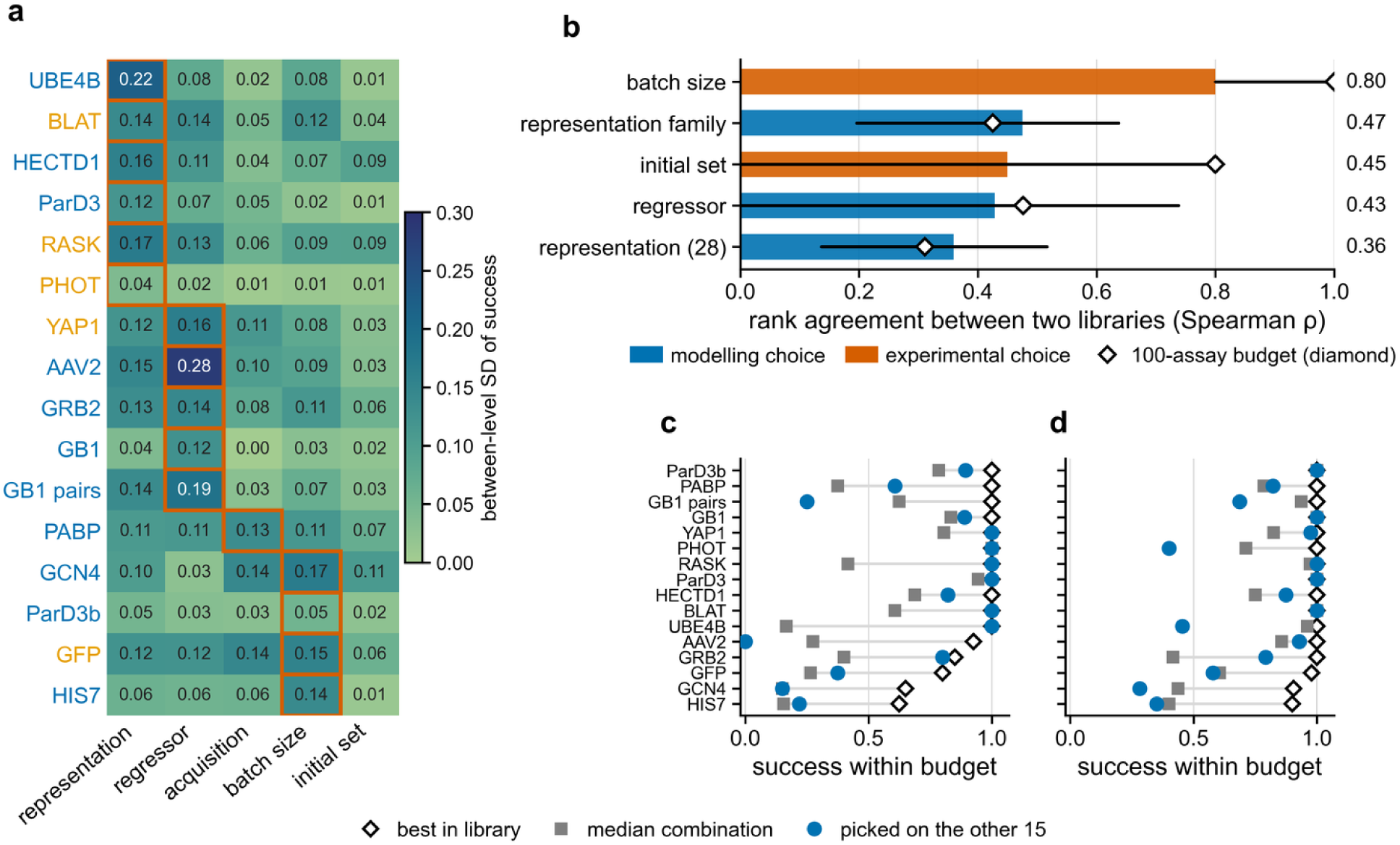
The importance and ranking of MLDE design choices vary between protein landscapes. (a) Variation in campaign success attributable to each design choice within individual landscapes after 400 newly acquired assays. Outlines indicate the most influential design choice for each landscape. Label colors distinguish combinatorial libraries (blue) and substitution scans (orange). (b) Agreement between landscape pairs in their rankings of the different levels of each design factor, expressed as Spearman rank correlations (median and interquartile range) after 400 acquired assays. Diamonds indicate the corresponding results after 100 acquisitions. (c,d) Success rates of representation–regression-model combinations selected using 15 landscapes and evaluated on the held-out sixteenth landscape, compared with the best-performing and median combinations on that landscape after 100 and 400 acquired assays, respectively. The “median combination” refers to the median success rate among all 224 evaluated combinations, rather than a separately optimized model.

**Figure 12.**
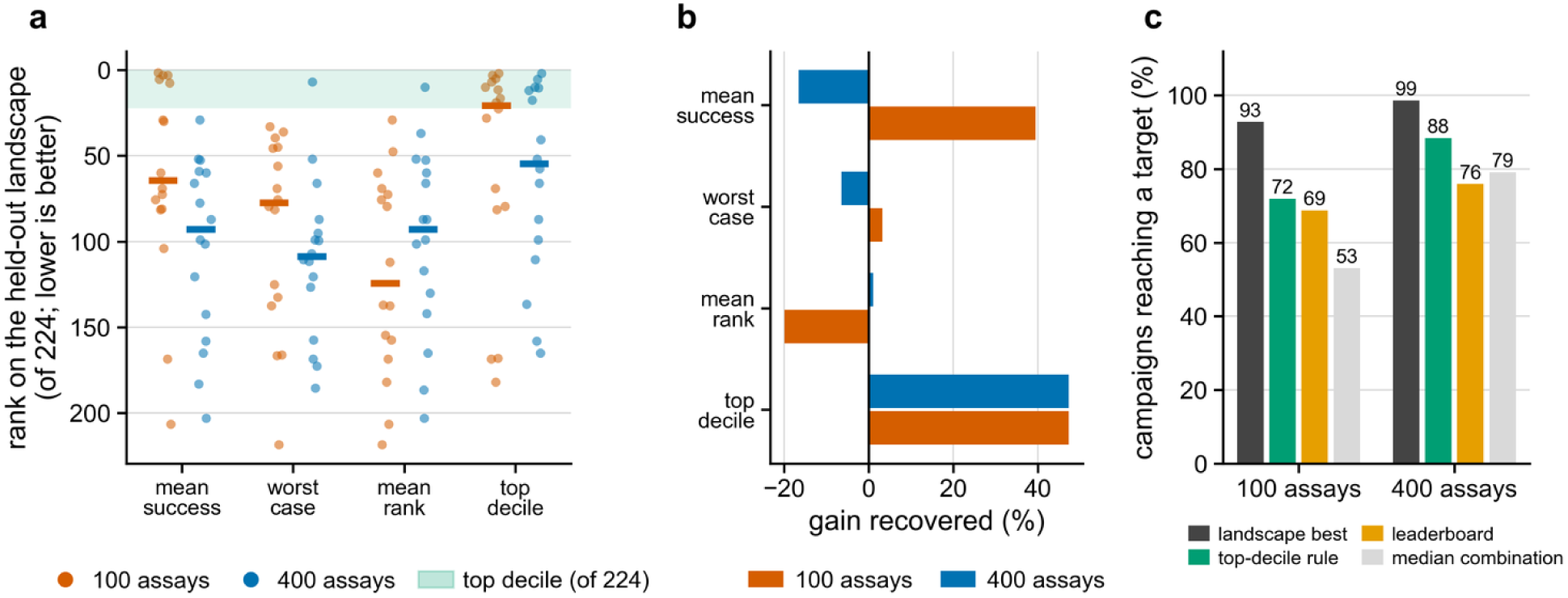
Comparison of four strategies for selecting an MLDE configuration for a previously unseen landscape. For each of the 16 evaluations, one landscape was held out, while a representation–regression-model combination was selected using the remaining 15 landscapes. Four selection criteria were compared: mean success, worst-case success, mean rank, and frequency of ranking among the top 10%. (a) Rank of each selected combination among all 224 combinations on the held-out landscape. Lower ranks indicate better performance, and the shaded region marks the top 10%. Points represent individual held-out landscapes, and horizontal bars indicate median ranks. (b) Fraction of the achievable improvement over the median-performing combination recovered by each selection criterion, relative to an idealized selection of the best combination separately for each landscape. Bars correspond, from top to bottom, to the four criteria shown in panel (a); negative values indicate performance below the median combination. Orange and blue indicate results after 100 and 400 acquired assays, respectively. (c) Success rates of the idealized landscape-specific selection, top-decile-frequency selection, mean-success selection, and the median combination after 100 and 400 acquired assays. Apparent differences between selection criteria should be interpreted in light of their statistical uncertainty.

The influence of individual design choices on successful variant identification also varied substantially between proteins. For every design choice, formal statistical tests rejected the assumption of a consistent effect across all 16 landscapes (all 40 comparisons, *p* ≤ 0.0005). For example, the I² statistic, which quantifies the proportion of observed variation in an effect attributable to differences between landscapes rather than sampling uncertainty, was 95.7% for representation choice (95% CI, 95.4–95.9%), 95.4% for regression model, and 96.4% for acquisition rule. Across the 16 landscapes, 13 different rankings of the five design factors according to their influence on campaign success were observed (Kendall’s W = 0.42^32^; **Figure 11a**). These results demonstrate that a single ranking of MLDE configurations can obscure substantial differences between protein fitness landscapes.

To investigate whether differences in MLDE performance between landscapes could be explained by measurable landscape properties, ten characteristics, including epistasis, navigability, library size, and the frequency of high-fitness variants, were examined for associations with campaign success and the effects of different MLDE design choices (**Method S6**, **Result S3**, **Tables S5** and **S6**). However, only a weak association was observed between the fraction of variants reaching at least 95% of the maximum fitness and the difficulty of identifying successful variants (Spearman ρ = 0.36, *p* = 0.17). None of the 100 tested associations remained statistically significant after Holm correction (lowest adjusted *p* = 0.38).

### Alternative ways to choose models do not reliably improve transfer

Finally, we examined whether the limited transferability of the highest-ranked configuration could be improved by applying alternative criteria for selecting representation–regression-model combinations from benchmark results. As before, 15 landscapes were used for selection and the remaining landscape for evaluation, with the procedure repeated for all 16 landscapes. Four selection criteria were compared: the highest average success (“mean success”); the highest success on the least favorable training landscape (“worst-case success”); the lowest average rank across training landscapes (“mean rank”); and the largest number of training landscapes on which a combination ranked among the top 10% (“top-decile frequency”). These criteria favor high average performance, avoidance of poor outcomes, consistently high relative rankings, and frequent placement among the best-performing combinations, respectively (**Figure 12a,b**).

Selection based on top-decile frequency yielded the highest mean success rate on held-out landscapes, reaching 88.4% after 400 acquired assays, compared with 75.9% for selection based on mean success (**Figure 12c**). However, this difference was not statistically supported by the uncertainty analysis and therefore does not establish a reliable improvement in transfer performance. When evaluated across all 16 landscapes rather than only the held-out landscapes, combinations selected by mean success achieved an average success rate of 89.2%, compared with 88.4% for the top-decile-frequency criterion. The highest-performing individual combination in the full benchmark, ProstT5 with random forest, achieved 89.7% success. Neither worst-case success nor mean rank provided a statistically supported improvement over selection by mean success, with reported differences of 2.0 and 3.4 pp (bootstrap intervals, −14.5 to 8.6 and −8.7 to 7.9, respectively).

The apparent advantage of top-decile frequency was largely explained by the fact that this criterion selected the same representation–model combination in all 16 leave-one-landscape-out evaluations. Consequently, the same average success rate was obtained when evaluated across all landscapes or only the held-out landscapes. In contrast, the mean-success criterion selected different combinations depending on which landscape was excluded, resulting in greater variation in held-out performance. To assess how much of this difference could arise from selecting the highest-scoring configuration on a finite benchmark, the selection procedure was repeated after randomly shuffling configuration identities within each landscape. Under this control, mean-success and top-decile-frequency selection yielded success rates of 71.6% and 73.7%, respectively, both below the 79.1% success rate of the median combination. These results demonstrate that configurations selected for their strong performance on the training landscapes may perform poorly on unseen landscapes, even when consistent performance differences between configurations have been removed.

Across the four selection criteria, the average reduction in success rate from the landscapes used for configuration selection to the held-out landscape was 13.3 pp (95% landscape-bootstrap interval, 1.1–26.5). It was further examined whether this reduction could be attributed to related datasets being included in both the selection and evaluation sets. When landscapes derived from the same protein were grouped and withheld together, a reduction of 15.7 pp was observed. Grouping landscapes originating from the same study yielded a reduction of 13.0 pp, compared with 13.3 pp for individual landscape hold-outs (**Figure S11**). Thus, the limited transferability of selected configurations persisted even when related datasets were excluded from the training portion of the benchmark.

Finally, the possibility of selecting a configuration based on information obtained during the first model-update round of an individual campaign was investigated. A fully prospective evaluation of this approach would require additional simulations. Therefore, an optimistic scenario was constructed in which the selection procedure had access to the accuracy with which every configuration ranked variants during the first round, although the necessary experimental fitness measurements would not all be available in practice. Despite this advantage, campaign-specific configuration selection did not outperform a single fixed configuration, with success rates of 68.8% versus 71.9% after 100 acquired assays and 88.2% versus 88.4% after 400 assays. Higher success rates with campaign-specific selection were observed on only three of the 16 landscapes after 400 assays. Moreover, early ranking accuracy was only moderately correlated with final campaign success (median Spearman ρ = 0.30 across 541 scenarios). These findings indicate that reliably selecting model configurations from limited early campaign data remains challenging.

## DISCUSSION

Three main conclusions can be drawn from our MLDEx benchmark of simulated directed-evolution campaigns across 16 experimental protein fitness landscapes. First, ML models trained on experimental measurements improved the efficiency of identifying high-fitness variants compared with random screening, particularly when initial training sets were small. However, this advantage was reduced when the experimental measurements required to establish the initial training sets were included in the assay budget. Moreover, zero-shot screening achieved comparable performance under the uniform-initialization protocol used here. Second, comparatively consistent effects of acquisition strategy and batch size were observed across landscapes. Third, the relative performance of protein representations and regression models varied substantially between landscapes, such that combinations ranked highly on existing proteins did not reliably perform well on previously unseen ones.

The findings regarding acquisition strategy and batch size have practical implications for MLDE campaign design. Greedy acquisition outperformed expected improvement on 14 of the 16 landscapes, suggesting that selecting variants with the highest predicted fitness provides a strong baseline for the tested models and uncertainty estimates. Smaller batches were advantageous when only a limited number of additional variants could be measured, as more frequent model updates were possible. However, in the additional experiment without early stopping, this advantage was reversed after approximately 220 acquisitions. Batch size should therefore be selected according to the planned experimental budget and the benefit of repeated model updates, rather than assuming that either small or large batches are universally superior.

The limited transferability of representation–model rankings reflects a broader challenge in machine-learning benchmarking, where the best-performing method can depend strongly on the datasets included in the comparison.^4,31^ In the present benchmark, configuration rankings were relatively consistent within individual fitness landscapes but showed considerably less agreement between proteins. Consequently, a representation–model combination selected using 15 landscapes was unlikely to rank among the best on the held-out sixteenth landscape. However, the observed reduction in held-out performance cannot be attributed entirely to biological differences between landscapes. A performance reduction was also observed after configuration identities were randomly shuffled within landscapes, demonstrating that selecting the highest-performing configuration from a finite benchmark can itself produce overly optimistic expectations. Both genuine landscape dependence and this statistical selection effect must therefore be considered when interpreting benchmark rankings.

No simple landscape descriptor was identified that reliably predicted which MLDE configuration would perform best. In previous work, landscape navigability was reported to be informative when comparing the benefits of ML guidance across combinatorial libraries.^14^ In contrast, none of the ten landscape descriptors examined here showed a significant association with configuration effects after correction for 100 comparisons. Several factors may explain this difference. The present benchmark included substitution scans in addition to combinatorial libraries, and some graph-based landscape descriptors cannot be reliably calculated for sparsely sampled landscapes. Furthermore, the relative effects of individual configuration choices were evaluated rather than the overall benefit of ML guidance. Finally, the number of landscapes was limited to 16, reducing the statistical power to detect associations after correction for multiple testing. The absence of significant associations should therefore not be interpreted as evidence that landscape properties are unimportant.

The comparison with zero-shot screening highlights a promising opportunity for improving MLDE initialization. In the present simulations, initial training variants were sampled uniformly, although pretrained protein models could have been used to rank variants before experimental fitness measurements became available. Previous studies have demonstrated that zero-shot or otherwise informed initial-set selection can improve ML-guided directed evolution.^3,14^ This observation is extended by the present finding that direct zero-shot screening achieved performance comparable to iterative ML-guided campaigns when the same total number of experimental assays was considered. However, because campaigns initialized using zero-shot scores were not simulated, it remains unclear whether combining informed initialization with iterative model updates would outperform either approach alone.

### Limitations

Several limitations should be considered when generalizing these findings. In the main simulations, the effects of batch size and the ten-round early-stopping rule were partly confounded. After 200 acquisitions, 36% of campaigns using batches of 10 had terminated without success, whereas none of those using batches of 100 had stopped for this reason. To address this limitation, an additional experiment without early stopping was performed. However, this experiment was restricted to an initial training-set size of 100 variants, four representations, and three regression models selected to cover a range of computational costs. The crossover at approximately 220 newly acquired assays, beyond which batches of 100 variants achieved higher success rates than batches of 10, was therefore established only for these settings and may not apply to all MLDE campaigns.

The reasons why particular MLDE configurations perform well on some protein fitness landscapes but poorly on others also remain unclear. With only 16 landscapes, the present benchmark may have had insufficient statistical power to identify associations between landscape properties and campaign performance (**Result S3**). More diverse fitness landscapes and additional descriptors may therefore be needed to predict which MLDE configurations are most effective for a given protein.

The simulations were based on previously measured fitness landscapes rather than prospective rounds of protein synthesis and experimental testing. Consequently, experimental noise and biases were inherited from the original assays, and variants absent from the measured landscapes could not be explored. Furthermore, the landscapes were not entirely independent. GB1 and GB1 pairs represent the same protein domain.^22,24^ The two ParD3 datasets also share a protein.^17,18^ Both stability datasets (UBE4B and HECTD1) originated from the same study.^16^ When related datasets were withheld together, the reduction in held-out performance still persisted. A reduction of 15.7 pp was observed when landscapes from the same protein were grouped, and 13.0 pp when datasets from the same study were additionally grouped (**Figure S11**). Nevertheless, the benchmark was limited to 16 landscapes and therefore does not provide a comprehensive assessment of transferability across unrelated protein families or experimental conditions.

Finally, the number of landscapes eligible for configuration comparisons varied with initial training-set size. Campaigns that already contained a top-0.1% variant before model training were excluded from analysis. Because this occurred more frequently with larger initial training sets, no eligible campaigns remained for some landscapes. Consequently, all 16 landscapes were included for initial training sets of 100 and 200 variants, but only 14 and 11 landscapes for initial sets of 500 and 1,000 variants, respectively. Differences in performance between initial training-set sizes may therefore partly reflect changes in the landscapes included in the analysis, rather than the effect of training-set size alone.

### Future directions

Future MLDE approaches could benefit from adapting campaign decisions as experimental data accumulate, rather than fixing the representation, regression model, acquisition rule, and batch size at the outset. For example, campaigns could be initialized with small batches and frequent model updates, followed by larger batches once predictions become more informative or experimental throughput becomes the limiting factor. Such adaptive strategies should be evaluated prospectively and on previously unseen fitness landscapes.

Another priority is the development of methods for predicting which MLDE strategies are most suitable for a particular protein. Sequence and structural descriptors, evolutionary information, and early experimental measurements could be combined to characterize the underlying fitness landscape. Benchmark analyses could thereby be extended beyond ranking methods by their average success toward identifying the conditions under which specific approaches are most effective.

Finally, both experimental and computational costs should be considered when designing practical MLDE workflows. In the present benchmark, substantial differences in model-fitting time were observed between configurations with comparable success rates, while acquisition batch size affected both the frequency of model updates and the number of experimental assays required. Joint optimization of computational resources, initial training-set size, and subsequent experimental measurements could therefore further improve the efficiency of ML-guided protein engineering.

## CONCLUSIONS

MLDEx provides a systematic evaluation of machine-learning-guided directed evolution based on 557,143 simulated campaigns across 16 experimentally measured protein fitness landscapes. With small initial training sets, ML guidance improved the efficiency of identifying high-fitness variants compared with random screening, but this advantage diminished as the initial training-set size increased. When initial training measurements were included in the total assay budget, zero-shot screening was also found to be competitive under the tested conditions.

Several broadly applicable principles for MLDE campaign design were identified. Greedy acquisition generally outperformed expected improvement, while smaller batches and more frequent model updates were beneficial early in a campaign. An additional experiment demonstrated that larger batches became advantageous as the number of acquired variants increased. In contrast, the relative performance of protein representations and regression models varied substantially between landscapes. No combination consistently outperformed all others across the 16 landscapes, and configurations selected using existing datasets did not reliably achieve superior performance on previously unseen landscapes. For configurations with comparable success rates, substantial differences in computational cost provided an additional criterion for model selection.

These findings demonstrate the value of systematic MLDE benchmarking for identifying broadly applicable design principles, evaluating experimental efficiency, and assessing computational trade-offs. However, rankings of individual prediction models should be interpreted in the context of the protein fitness landscapes used for their evaluation rather than assumed to generalize to previously uncharacterized proteins.

## METHODS

### Protein fitness landscapes

We selected 16 experimentally measured deep mutational scanning (DMS) datasets from ProteinGym release 1.3.^7,33^ The datasets comprise 11 combinatorial libraries and five substitution scans, cover four assay classes (binding, activity, stability, and organismal fitness), and contain between 2,638 and 536,962 measured variants (**Table 1**). ProteinGym accession identifiers, protein lengths, and library types are listed in **Table S1**. We used the reported DMS fitness scores and normalized them separately for each landscape to a range of 0–1. This monotonic transformation preserves variant rankings and therefore does not affect rank-based analyses or the definition of top-performing variants.

### Simulation of directed-evolution campaigns

Each simulated campaign began with **n_start_** experimentally characterized variants sampled from a given fitness landscape. The selected representation and regression model were used to train a fitness predictor. The model then scored previously unmeasured **v**ariants, and the acquisition strategy selected **n_gain_** candidates for the next round. Their experimentally measured fitness values were retrieved from the landscape, added to the training set, and used to update the model. We simulated up to 40 acquisition rounds. Campaigns were terminated earlier if a variant reached at least 95% of the maximum measured fitness in that landscape or if ten consecutive rounds produced no improvement. The latter condition was defined as the early-stopping rule.

Initial training sets were generated using reproducible random seeds, and no predictor-based prefiltering was applied to the remaining candidate pool. We evaluated 28 sequence representations (**Table S2**): one-hot encoding; BLOSUM45, BLOSUM50, BLOSUM62, BLOSUM80, and BLOSUM90^34^; the Georgiev,^35^ Kidera,^36^ and physicochemical descriptor sets^37^; the sequence-based protein language models ESM-2,^38^ ProtT5,^39^ Ankh-base,^40^ and ESM-C,^41^ together with the structure-aware ProstT5^42^; SaProt,^43^ ProSST,^44^ VenusREM,^45^ ESM-IF1,^30^ ProteinMPNN probabilities,^46^ and SLAE^47^; and eight variants of these representations restricted to mutated positions or calculated as differences from the wild-type sequence. These groups cover simple residue descriptors, language-model embeddings, structure-informed features, and mutation-focused encodings. For ESM-C, the readout layer was selected separately for each landscape: at each of the 36 candidate layers, residue embeddings of 2,000 randomly sampled experimentally measured fitness variants were mean-pooled and a ridge regression probe^48^ was fitted under three-fold cross-validation, and the layer whose held-out predictions showed the highest Spearman correlation with measured fitness was retained. This landscape-specific selection step used experimental fitness information and was not applied to the other encoders **(Method S1** and **Table S3)**.

We combined each representation with eight regression models: ordinary linear regression, ridge regression,^48^ elastic net,^49^ support vector regression (SVR),^50^ random forest,^51^ LightGBM,^52^ XGBoost,^53^ and a multilayer perceptron (MLP). See **Table S4** for the regression models and software references.^54^ A ninth model, Gaussian-process regression,^55^ was evaluated but excluded from the final comparisons because completed runs were unevenly distributed across the landscapes **(Method S2** and **Table S4)**. Three candidate-selection strategies were evaluated: greedy acquisition (selecting variants with the highest predicted fitness), expected improvement^56^ (prioritizing variants predicted to improve upon the best measured fitness while accounting for prediction uncertainty), and uniform random acquisition as a control. For expected improvement, predictive uncertainty was estimated from the Gaussian-process posterior, variability among random-forest trees, or an ensemble of up to ten bootstrap-trained models for the other regressors. Results for expected improvement therefore reflect this particular implementation.

We tested four initial training-set sizes (**n_start_** = 100, 200, 500, or 1,000 variants), four acquisition batch sizes (**n_gain_**= 10, 20, 50, or 100 variants per round), and three replicate random seeds. Model hyperparameters were optimized using Optuna every five acquisition rounds.^57^ At least 20 optimization trials were performed, with a minimum of ten trials per tuned hyperparameter (**Table S4**). Optimization maximized the mean Spearman correlation between predicted and measured fitness using three-fold cross-validation on the labeled training set. Although the number of optimization trials was adjusted according to the number of tuned parameters, this does not guarantee equally effective hyperparameter optimization across models.

We recorded the first round in which a successful variant was identified and evaluated campaign success after specified numbers of newly acquired variants (**A** = 100 or 400 unless otherwise stated; **Method S3**). These counts exclude the initial training set. Campaigns that stopped without finding a successful variant before reaching the specified number of acquisitions were scored as unsuccessful. For comparisons that also include the initial training set, the total number of experimental assays was calculated as **T** = **n_start_** + **A**.

### Definition of successful variants and analyzed campaigns

A successful campaign was defined as one that identified at least one variant among the top 0.1% of the measured fitness landscape (primary success criterion). This percentile-based criterion allows comparisons between libraries with different fitness distributions. We also used a separate stopping criterion based on identifying a variant with a fitness of at least 95% of the maximum measured value. However, this threshold was not used as the primary success criterion because only one or two variants qualified in 12 of the 16 landscapes. When fitness scores were tied at the percentile threshold, all tied variants were considered successful. The primary criterion was evaluated at specified numbers of assays rather than at campaign termination, allowing campaigns with different batch sizes to be compared at matched assay counts.

The complete benchmark comprised 557,143 completed campaigns. We excluded 41,075 Gaussian-process campaigns because they were not evenly represented across landscapes, leaving 516,068 campaigns for the principal analyses. Uniform sampling of the initial training set excluded variants at or above the 95%-of-maximum stopping threshold, but did not necessarily exclude all variants among the top 0.1% of fitness scores. Consequently, 29.2% of non-Gaussian-process runs already contained a successful variant before model training (**Method S4; Figure S1**). These runs were excluded from comparisons between model configurations, leaving 365,563 campaigns, of which 243,708 used ML-guided acquisition. Because all configurations within a scenario shared the same starting set, this exclusion did not favor any particular configuration. For comparisons with random or zero-shot screening, we retained campaigns with an initially successful variant because their initial measurements contributed to the total assay count. In two landscapes, reaching the 95%-of-maximum stopping criterion could terminate a campaign before identification of a variant satisfying the more stringent top-0.1% criterion; these runs were treated as right-censored where appropriate. An additional 28 planned runs were not completed because of computational cost (less than 0.01% of planned campaigns).

### Statistical analysis

Campaigns with the same landscape, initial-set size, acquisition batch size, and random seed shared their initial variants. We therefore treated this combination, termed a scenario, as the unit for resampling and confidence-interval estimation. To prevent large landscapes or landscapes with more completed campaigns from dominating comparisons, summary statistics were calculated first within each landscape and then averaged across the 16 landscapes with equal weight (landscape-balanced means).

To determine whether the effect of each design choice differed between landscapes, we fitted generalized estimating equations^58^ separately for each design factor and landscape, using an independence working correlation and CR2-corrected cluster-robust standard errors.^59,60^ Variation in the estimated effects across landscapes was quantified with Cochran’s *Q* and *I²* statistics and Higgins–Thompson confidence intervals.^61,62^ The eight specifications shown in **Figure S2** used logit and identity links at the 100- and 400-assay endpoints and included either all acquisition arms or ML-guided arms only (**Method S5**). **Method S4** describes the treatment of campaigns that succeeded during initialization. For key comparisons, we also used a bootstrap resampling complete scenario, an exact sign-flip test of the 16 landscape-level differences, and a wild-cluster bootstrap with Rademacher weights.^63,64^ Associations between ten fitness-landscape descriptors (including epistasis, navigability, and target density; **Method S6** and **Table S5**) and design-factor effects were tested in 100 comparisons, with Holm correction for multiple testing.^65^

### Performance measures

Success was defined as the proportion of campaigns that identified a variant within the top 0.1% of the fitness landscape by a specified assay count. We distinguish **A**, the number of variants tested after model training began, from **T**, the total number of experimental measurements, including the initial training set (**T** = **n_start_** + **A**). Campaigns that could not reach a specified total assay count within the 40-round limit were excluded from the corresponding comparison rather than classified as failures. Success rates at fixed numbers of acquired assays were analyzed separately, as described above.

Landscape-balanced success was calculated as the unweighted mean of the success rates across individual landscapes. Thus, each of the 16 landscapes contributed equally, regardless of the number of simulated campaigns.

We evaluated the transfer of model configurations to previously unseen landscapes using a leave-one-landscape-out approach. For each iteration, we selected a representation–model combination based on its performance across 15 landscapes and evaluated its success on the remaining landscape. The transfer gap was defined as the difference between the average performance of the selected combinations on the landscapes used for selection and their average performance on the held-out landscapes. Because the selected combination could differ between iterations, this measure does not correspond to the difference between the training and held-out performance of a single globally selected configuration.

The fraction of achievable improvement recovered by a selection strategy was calculated as (s_rule_ – s_median_) / (s_oracle_ − s_median_), where s_rule_ is the success rate of the selected combination, s_median_ is the median success rate across all tested combinations, and s_oracle_ is the average success rate obtained by selecting the best combination separately for each landscape. A value of 100% represents this idealized landscape-specific selection, whereas negative values indicate performance below that of the median combination.

### Screening controls and additional experiments

#### Random screening

To estimate the success probability of uniformly screening variants without ML guidance, we used the analytical expression P(success after T assays) = 1 − (1 − p)^T^, where p is the proportion of successful variants in each landscape. We used this analytical baseline rather than simulated random-acquisition campaigns because the latter were subject to the same round-based early-stopping rule as ML-guided campaigns, which could bias comparisons with unrestricted random screening.

#### Zero-shot screening

As an alternative that does not require experimental fitness measurements for ranking, variants were scored using the pretrained structure-conditioned models ESM-IF1^30^ and ProteinMPNN.^46^ Each variant was assigned a score corresponding to the average log-probability of its amino acids at the mutated positions. For a total budget of **T** assays, the **T** highest-ranked variants were selected directly. Scores were obtained from previously computed per-position probability matrices, without model retraining or the use of experimental fitness labels. Results for ESM-IF1 are presented in the main text, whereas those for ProteinMPNN are provided in **Result S1**. Variants lacking a valid score at any mutated position were excluded, and ties were resolved according to library order.

#### Permutation control for configuration selection

To determine whether apparent performance losses could arise from the configuration-selection procedure itself, we randomly shuffled configuration identities within each landscape and repeated the selection and evaluation procedure. This preserved differences in difficulty between landscapes while removing consistent performance advantages associated with individual configurations.

#### Bootstrap for landscape transfer

We resampled the landscapes with replacement and repeated the complete leave-one-landscape-out procedure to estimate the uncertainty in configuration performance on previously unseen landscapes.

*Within-campaign model selection.* We also evaluated an optimistic scenario in which a campaign could select its representation–model combination based on the first-round ranking results from all combinations. This approach requires experimental fitness information that would not normally be available at the time of model selection. It therefore represents an upper bound for this early-data selection strategy rather than a directly applicable prospective approach.

#### Experiment without early stopping

To distinguish the effect of batch size from that of the early-stopping rule, which terminates campaigns after ten consecutive rounds without improvement (**n_cancel_** = 10), we performed 4,608 additional simulations with **n_cancel_** = 40. This setting prevented termination due to a lack of improvement before acquiring 400 variants. Each campaign was designed to acquire exactly 400 variants, regardless of batch size. The experiment included all 16 landscapes, an initial training set of 100 variants, four acquisition batch sizes (10, 20, 50, and 100), two acquisition rules (greedy and expected improvement), three random seeds, and four representations combined with three regression models selected to cover a range of computational costs. Of the 4,608 planned campaigns, 4,593 were completed, and 4,497 remained eligible after excluding 96 campaigns that succeeded during initialization. These results were analyzed separately from the main benchmark.

#### Variance decomposition

For each landscape, we partitioned the variation in success across all 28 × 8 representation–regression-model combinations into contributions from the representation, the regression model, and their interaction. The proportions of total variation attributable to these three components were averaged across landscapes, and confidence intervals were estimated by resampling landscapes with replacement.

## Abbreviations

MLDE: machine-learning-guided directed evolution
DMS: deep mutational scanning
PLM: protein language model
EI: expected improvement
SVR: support vector regression
RF: random forest
MLP: multilayer perceptron
GP: Gaussian process
CI: confidence interval
GEE: generalized estimating equations
CR2: bias-reduced linearization
CPU: central processing unit.

Further details regarding the methods can be found in the supplemental information.

## RESOURCE AVAILABILITY

### Lead contact

Requests for information and resources should be directed to Georg Künze.

### Materials availability

This computational study used previously published datasets and generated no new physical materials.

### Data and code availability

- The complete run log (MLDEx, 557,143 campaigns), the derived tables behind every figure, and the analysis plan are archived at https://doi.org/10.5281/zenodo.23320790 under CC BY 4.0. Source landscapes are from ProteinGym release 1.3^33^ (archive DMS_ProteinGym_substitutions.zip, MD5 ca1a4d46941ef33cc972245347118c7d, retrieved 2026-08-10).
- The simulation and analysis code is available at https://github.com/kuenzelab/MLDEx under the MIT license.
- Any additional information required to reanalyze the data reported in this paper is available from the lead contact upon request.

## ACKNOWLEDGMENTS

Research in the Künze laboratory is supported by the Deutsche Forschungsgemeinschaft (DFG) through project 421152132 (CRC 1423, subproject C07), project 448298270 (TRR 386, subprojects A2 and B2), and project 514901783 (CRC 1664, subproject D01), and by the Sächsisches Staatsministerium für Wissenschaft, Kultur und Tourismus (SMWK; project 100704504). The authors further acknowledge financial support from the German Federal Ministry of Education and Research and the SMWK through the Center of Excellence for AI Research “Center for Scalable Data Analytics and Artificial Intelligence Dresden/Leipzig” (project identification number: ScaDS.AI). We thank the Leipzig University Computing Centre and the Centre for Information Services and High Performance Computing (ZIH) at TU Dresden for computational resources.

## AUTHOR CONTRIBUTIONS

G.K., conceptualization, supervision, and funding acquisition; A.T., investigation, data curation, and initial analysis; R.S., software, methodology, formal analysis, data curation, visualization, and writing original draft; R.S. and G.K., writing review and editing. All authors approved the final version of the manuscript.

## DECLARATION OF GENERATIVE AI AND AI-ASSISTED TECHNOLOGIES IN THE WRITING PROCESS

During the preparation of this manuscript, the authors used OpenAI to assist with manuscript formatting, and editing the Summary and textual refinement. After using this tool, the authors reviewed and edited the manuscript as needed and take full responsibility for the content of the publication.

## SUPPLEMENTAL INFORMATION

Document S1. Accessions for the 16 landscapes; the representation, regression-model and ESM-C read-out tables; the analyzed population; per-landscape results, budget curves and the landscape covariate analysis (Methods S1–S6, Results S1–S3, Figures S1–S11, Tables S1–S6).

