## Supporting Information for "Evaluating the Generalizability of Machine-Learning-Guided Directed Evolution Across Protein Fitness Landscapes"

### Supporting Methods

**Table S1** identifies the experimental datasets from ProteinGym release 1.3.[1] **Table S2** lists the representations. The methods below describe model and campaign eligibility, endpoint evaluation, and the analysis of differences between landscapes.

***Method S1. Landscape-specific selection of the ESM-C readout layer***

ESM-C[2] was included as a protein representation in every analysis. Unlike the other representations, which used their standard readout, the ESM-C readout layer was selected separately for each landscape based on experimental fitness data from a probe set of 2,000 variants (**Table S3**). This additional layer-selection step should be considered when comparing ESM-C with other representations.

***Method S2. Exclusion of Gaussian-process regression from the main analyses***

A total of 41,075 campaigns using Gaussian-process regression[3] were completed, but their coverage across landscapes was uneven because of the model's computational cost. These campaigns were therefore excluded from the main analyses rather than being compared using an incomplete set of simulations. The other regression models retained data in all 3,584 configuration-factor cells of the analyzed design (**Table S4**). For any pooled marginal estimate, the maximum possible change caused by omitting Gaussian-process runs is bounded by the fraction of runs removed from that estimate.

***Method S3. Evaluation at matched numbers of experimental assays***

The maximum number of acquisition rounds was fixed, so campaigns with larger batches could test more variants before termination. Comparing success only at termination would therefore confound the effect of batch size with the number of experiments performed. To make comparisons fair, assay-dependent outcomes were evaluated at fixed numbers of newly acquired variants, and the number of assays needed to identify the first successful variant was reported separately. In two landscapes, a campaign could terminate after reaching the secondary stopping threshold (95% of maximum fitness) without meeting the primary top-0.1% success criterion. Such campaigns were treated as right-censored at termination and excluded from the relevant denominator.

***Method S4. Exclusion of campaigns already successful at initialization***

Initial training sets were sampled uniformly from each landscape after excluding variants with fitness at or above the secondary stopping threshold (95% of the landscape maximum). However, this sampling rule did not exclude all variants in the top 0.1% of the fitness landscape, which defined the primary success criterion. In most landscapes, the primary threshold was lower than the secondary stopping threshold. For a target density d (the fraction of variants meeting the primary success criterion), the probability that an initial sample of $n_{start}$ variants already contains a target is $1 - (1 - d)^{n_{start}}$. This occurred in 29.2% of the non-Gaussian-process campaigns, with the frequency increasing with initial training-set size as predicted by the analytical expectation (**Figure S1**, manuscript **Figure 1**, and E3_start_contamination.csv). Such campaigns had already succeeded before a prediction model was fitted and therefore provided no information about the effectiveness of an MLDE configuration.

These campaigns were excluded from comparisons between configurations, leaving 365,563 eligible campaigns, including 243,708 with ML-guided acquisition. All acquisition arms within a scenario used the same initial training set, so removing campaigns that had already succeeded did not favor any configuration; retaining them would instead reduce the apparent differences between arms. Success rates reported for configuration comparisons are therefore conditional on an initial set that contained no primary target. By contrast, comparisons with random and zero-shot screening retained the initial measurements and all initially successful campaigns. Their absolute success rates should therefore not be directly compared with those from the configuration analyses.

***Method S5. Estimation of differences between protein landscapes***

Differences in the effects of MLDE design choices between landscapes were analyzed using generalized estimating equations with logit and identity links.[4] The identity link estimates differences in success probabilities directly; these risk differences are collapsible and are less affected by near-complete separation when success rates approach 100%, which can make logit estimates unstable. All eight analysis specifications were reported: two assay budgets (100 and 400 newly acquired variants), two link functions (logit and identity), and two sets of campaigns (all acquisition arms or ML-guided arms only; **Figure S2**; T2_heterogeneity_cr2.csv). One of the eight models, for PHOT with a logit link, ML-guided arms only, and the 400-assay endpoint, did not produce plausible converged coefficient estimates and was flagged in T2_gee_fit_diagnostics.csv. The corresponding identity-link analysis was unaffected, and no reported conclusion depends on the problematic fit. Heterogeneity was evaluated with Cochran's Q and $I^2$ and Higgins–Thompson confidence intervals.[5,6] Applying the CR2 correction[7,8] increased the standard errors by a median factor of 1.039; the highest cluster leverage was 0.840.

***Method S6. Calculation of the fitness landscape descriptors***

We calculated fitness landscape descriptors directly from experimentally measured fitness values, without using structural information or predictive models. Variants were considered neighbors if they differed by a single amino acid substitution. Library size was defined as the number of variants with valid mutation annotations, and target density as the number of variants with fitness values at least 95% of the maximum fitness per $10^5$ library variants. The assayed fraction represented the total campaign budget, including initial training and acquisition rounds, relative to library size, and could exceed 100% for small libraries. Fitness dynamic range was measured using the interquartile range (IQR), while mutational complexity was described by the fraction of variants with two or more substitutions and the number of distinct mutated positions. Rank epistasis was calculated as 1 − Spearman's ρ between measured multiple-mutant fitness and additive predictions based on constituent single mutants. This rank-based measure allowed comparisons across independently normalized ProteinGym assays. Epistasis was calculated only when at least 25% of multiple mutants had complete single-mutant measurements, with a minimum of 30 eligible variants. Landscape ruggedness was characterized by the number of local fitness optima per $10^5$ connected variants, where a local optimum had fitness equal to or greater than all measured neighbors. Navigability was defined as the fraction of connected variants with a strictly fitness-increasing single-mutation path to a target variant. Navigability was previously investigated as a factor influencing MLDE performance by Li et al.[9] Both metrics were excluded when fewer than 50% of variants had measured neighbors. Associations between landscape descriptors and ten campaign outcomes were evaluated using Spearman correlation, requiring at least six landscapes with valid measurements per comparison. The resulting family of 100 tests was corrected for multiple comparisons using the Holm procedure.[10] **Table S5** gives the descriptor values.

### Supporting Results

##### *Result S1: Zero-shot protein models*

Two pretrained, structure-conditioned protein models, ESM-IF1[11] and ProteinMPNN,[12] were evaluated for zero-shot screening. Each variant was scored using the mean log-probability assigned to its amino acids at the mutated positions. Scores from both models generally ranked variants in the same direction, but ESM-IF1 showed a stronger association with measured fitness. Across the 16 landscapes, the mean Spearman correlation between zero-shot score and experimental fitness was 0.37 for ESM-IF1 and 0.29 for ProteinMPNN. Values for individual landscapes are provided in zeroshot.csv. Because of its stronger correlation with measured fitness, ESM-IF1 was used for the zero-shot comparison in the main text.

##### *Result S2: Batch-size effects and the crossover in success rates*

**Figure S4** shows campaign success as the number of acquired assays increased for each acquisition batch size and landscape. **Figure S5** identifies the assay count at which batches of 100 variants began to outperform batches of 10 in success rate. Across the 16 landscapes where the success curves crossed, this crossover occurred between 104 and 280 newly acquired assays; the pooled estimate was 176. These values come from the main benchmark, in which the early-stopping rule was active. In the additional experiment without early stopping, the crossover occurred at approximately 220 acquired assays (main-text Section 3.3). In the main benchmark, the reversal beyond the first 100 acquisitions was partly confounded by the stopping rule, which was defined by the number of rounds rather than the number of tested variants (T5_early_stop.csv). The additional experiment confirmed that the reversal persisted when early stopping due to a lack of improvement was prevented.

##### *Result S3: Relationships between landscape properties and MLDE performance*

Ten characteristics of each protein fitness landscape were calculated from the experimental fitness tables: rank epistasis; number of local fitness optima per $10^5$ connected variants; navigability; fraction of connected variants; fitness dynamic range; fraction of variants containing multiple mutations; number of mutated positions; library size; target density (the fraction of target variants); and the fraction of the library that a campaign would assay. Their values for each landscape are given in **Table S5**, and the calculation of each is described in **Method S6**. Each was tested for association with the observed effect of MLDE design choices on campaign success, giving 100 comparisons in total (**Table S6**). None remained statistically significant after Holm correction for multiple testing[10] (smallest adjusted $p$ = 0.38). Rank epistasis was not assigned a value for substitution scans, where multiple mutations were present only in an incompletely sampled subset rather than in a deliberately constructed combinatorial library. Among the 11 combinatorial landscapes it was additionally undefined for GCN4 and ParD3b, where fewer than a quarter of the multiple-mutant variants had all of their constituent single mutants measured, leaving nine landscapes. Measures of landscape ruggedness and navigability were also left undefined when fewer than half of the measured variants had at least one measured sequence differing at a single amino-acid position (a Hamming-distance-1 neighbor), leaving 13 landscapes. Without this restriction, sparsely sampled landscapes could incorrectly appear maximally rugged.

#### Supporting Tables

**Table S1. ProteinGym accession identifiers and characteristics of the 16 protein fitness landscapes.** The identifiers are from ProteinGym release 1.3.[1] The year in an accession can differ from the final publication year. The landscape column cites the original studies.

| Landscape | ProteinGym DMS id | UniProt | Construction | Length | Variants |
|---|---|---|---|---|---|
| GFP[13] | GFP_AEQVI_Sarkisyan_2016 | GFP_AEQVI | substitution | 238 | 51,714 |
| YAP1[14] | YAP1_HUMAN_Araya_2012 | YAP1_HUMAN | substitution | 504 | 10,075 |
| RASK[15] | RASK_HUMAN_Weng_2022_binding-DARPin_K55 | RASK_HUMAN | substitution | 188 | 24,873 |
| PHOT[16] | PHOT_CHLRE_Chen_2023 | PHOT_CHLRE | substitution | 118 | 167,529 |
| GB1[17] | SPG1_STRSG_Wu_2016 | SPG1_STRSG | combinatorial | 448 | 149,360 |
| GB1 pairs[18] | SPG1_STRSG_Olson_2014 | SPG1_STRSG | combinatorial | 448 | 536,962 |
| ParD3[19] | F7YBW8_MESOW_Aakre_2015 | F7YBW8_MESOW | combinatorial | 93 | 9,192 |
| ParD3b[20] | F7YBW8_MESOW_Ding_2023 | F7YBW8_MESOW | combinatorial | 93 | 7,922 |
| GCN4[21] | GCN4_YEAST_Staller_2018 | GCN4_YEAST | combinatorial | 281 | 2,638 |
| HIS7[22] | HIS7_YEAST_Pokusaeva_2019 | HIS7_YEAST | combinatorial | 220 | 496,137 |
| GRB2[23] | GRB2_HUMAN_Faure_2021 | GRB2_HUMAN | combinatorial | 217 | 63,366 |
| AAV2[24] | CAPSD_AAV2S_Sinai_2021 | CAPSD_AAV2S | combinatorial | 735 | 42,328 |
| PABP[25] | PABP_YEAST_Melamed_2013 | PABP_YEAST | combinatorial | 577 | 37,708 |
| UBE4B[26] | UBE4B_HUMAN_Tsuboyama_2023_3L1X | UBE4B_HUMAN | combinatorial | 69 | 3,622 |
| HECTD1[26] | HECD1_HUMAN_Tsuboyama_2023_3DKM | HECD1_HUMAN | combinatorial | 72 | 5,586 |
| BLAT[27] | BLAT_ECOLX_Firnberg_2014 | BLAT_ECOLX | substitution | 286 | 4,783 |

The following datasets were requested but were not available in ProteinGym release 1.3: DHFR, T7-RNAP, TEV, TrpB1, TrpB3E, TrpB8, and PhoQ. No substitute datasets were used.

**Table S2. The 28 protein representations grouped by family.** The table lists the representation names used in the benchmark and the number of representations in each family.

| Family | Members | |
|---|---|---|
| one-hot | one_hot | 1 |
| substitution matrix | blosum45, blosum50, blosum62, blosum80, blosum90[28] | 5 |
| physicochemical | georgiev,[29] kidera,[30] physchem[31] | 3 |
| protein language model | esm2_t33_650M,[32] prot_t5,[33] prost_t5,[34] ankh_base,[35] esmc_600m[2] | 5 |
| structure-based | saprot_650m,[36] prosst,[37] venus_rem,[38] esm_if1,[11] proteinmpnn_probs,[12] slae[39] | 6 |
| site-restricted | site:georgiev, site:prot_t5, site:esm2_t33_650M, site:slae | 4 |
| delta (variant WT) | delta:georgiev, delta:prot_t5, delta:esm2_t33_650M, delta:slae | 4 |

**Table S3. ESM-C[2] readout layers selected for the individual fitness landscapes.** For each landscape, the selected layer gave the highest reported Spearman correlation (ρ) between the ridge probe's held-out predictions and measured fitness in a probe set of 2,000 variants from that landscape. The table compares this selected layer with the default final layer. Both correlation columns refer to the same probe set, which is separate from the landscape-level hold-out used to evaluate configuration transfer (main-text Section 2.5). Other representations were not subjected to an equivalent search over readout layers, and ESM-C results should be interpreted with that difference in mind.

| Landscape | Layer | Held-out | Final-layer |
|---|---:|---:|---:|
| AAV2 | 16 | 0.770 | 0.598 |
| BLAT | 33 | 0.752 | 0.639 |
| GB1 | 17 | 0.542 | 0.264 |
| GB1 pairs | 28 | 0.884 | 0.811 |
| GCN4 | 18 | 0.557 | 0.203 |
| GFP | 11 | 0.740 | 0.407 |
| GRB2 | 21 | 0.780 | 0.644 |
| HIS7 | 15 | 0.776 | 0.569 |
| PABP | 33 | 0.820 | 0.754 |
| ParD3 | 3 | 0.462 | 0.237 |
| ParD3b | 18 | 0.775 | 0.515 |
| PHOT | 15 | 0.953 | 0.879 |
| RASK | 25 | 0.883 | 0.674 |
| UBE4B | 15 | 0.875 | 0.644 |
| HECTD1 | 17 | 0.885 | 0.714 |
| YAP1 | 16 | 0.649 | 0.521 |

**Table S4. Regression models and allocation of hyperparameter-optimization trials.** Each model was allocated max(20, 10 × h) Optuna trials,[40] where h is the number of tuned hyperparameters. Thus, at least 20 trials were used, with more trials for models with more tuned parameters. Gaussian-process regression was discontinued because of its high computational cost and was excluded from the final comparisons. The software references include scikit-learn.[41]

| Model | Family | Hyperparameters | Optuna trials | Note |
|---|---|---:|---:|---|
| linear | linear | 0 | 20 | |
| ridge | linear | 1 | 20 | [42] |
| elastic_net | linear | 2 | 20 | [43] |
| svr | kernel | 3 | 30 | [44] |
| gaussian_process | kernel | 1 | 20 | dropped; native predictive variance[3] |
| rf | tree ensemble | 4 | 40 | native predictive variance[45] |
| lightgbm | tree ensemble | 5 | 50 | [46] |
| xgboost | tree ensemble | 6 | 60 | [47] |
| mlp | neural network | 2 | 20 | |

**Table S5. Values of the ten landscape descriptors for each of the 16 fitness landscapes.** Descriptors are defined in **Method S6**. Em dashes ("—") indicate descriptors that were left undefined: rank epistasis for substitution scans and for the two combinatorial landscapes with insufficient coverage of constituent single mutants, and the two graph-based measures for landscapes in which fewer than half of the variants had a measured Hamming-distance-1 neighbor.

| Landscape | Size | Density (per $10^5$) | Assayed (%) | Epistasis | Optima (per $10^5$) | Navigab. | Connected | Dyn. range | Multi-mut. | Positions |
|---|---|---|---|---|---|---|---|---|---|---|
| AAV2 | 42,328 | 4.7 | 11.81 | 0.717 | 12,859.2 | 0.564 | 0.716 | 5.269 | 0.987 | 28 |
| BLAT | 4,783 | 20.9 | 104.54 | — | 20.9 | 1.000 | 1.000 | 0.886 | 0.000 | 263 |
| GB1 | 149,360 | 0.7 | 3.35 | 0.807 | 123.9 | 0.997 | 1.000 | 0.008 | 0.999 | 4 |
| GB1 pairs | 536,962 | 1.9 | 0.93 | 0.042 | 1.7 | 1.000 | 1.000 | 4.569 | 0.998 | 55 |
| GCN4 | 2,638 | 37.9 | 189.54 | — | — | — | 0.070 | 0.342 | 0.987 | 44 |
| GFP | 51,714 | 150.8 | 9.67 | — | — | — | 0.423 | 2.187 | 0.979 | 233 |
| GRB2 | 63,366 | 3.2 | 7.89 | 0.138 | 127.8 | 0.940 | 1.000 | 0.749 | 0.984 | 56 |
| HECTD1 | 5,586 | 125.3 | 89.51 | 0.348 | 196.9 | 0.902 | 1.000 | 1.326 | 0.777 | 72 |
| HIS7 | 496,137 | 0.2 | 1.01 | 0.909 | 3,973.9 | 0.808 | 0.997 | 0.995 | 1.000 | 171 |
| PABP | 37,708 | 2.7 | 13.26 | 0.098 | 265.2 | 0.570 | 1.000 | 0.749 | 0.969 | 75 |
| PHOT | 167,529 | 1.2 | 2.98 | — | 46.6 | 0.999 | 1.000 | 0.452 | 0.987 | 118 |
| ParD3 | 9,192 | 21.8 | 54.40 | 1.157 | 761.5 | 0.985 | 1.000 | 0.013 | 0.996 | 4 |
| ParD3b | 7,922 | 12.6 | 63.12 | — | — | — | 0.371 | 0.411 | 0.990 | 10 |
| RASK | 24,873 | 12.1 | 20.10 | — | 44.2 | 0.954 | 1.000 | 0.760 | 0.876 | 187 |
| UBE4B | 3,622 | 27.6 | 138.05 | 0.336 | 497.0 | 0.791 | 1.000 | 1.252 | 0.691 | 69 |
| YAP1 | 10,075 | 19.9 | 49.63 | — | 585.6 | 0.816 | 1.000 | 0.872 | 0.964 | 34 |

**Table S6. All 100 tested associations between landscape descriptors and the effects of MLDE design choices.** Each row reports a Spearman correlation between one descriptor (**Table S5**) and one outcome, the number of landscapes for which both were defined, and the p value before and after Holm correction[10] across all 100 tests. Rows are ordered by uncorrected p value. No association remained significant after correction.

| Descriptor | Outcome | n | Spearman rho | p | p (Holm) |
|---|---|---|---|---|---|
| Dynamic range (IQR) | Effect of representation (SD) | 16 | 0.679 | 0.0038 | 0.379 |
| Multi-mutant fraction | Effect of representation (SD) | 16 | -0.671 | 0.0045 | 0.442 |
| Mutated positions | Batch 10 − 100 at 100 assays | 16 | -0.595 | 0.0151 | 1.000 |
| Mutated positions | Effect of batch size (SD) | 16 | 0.534 | 0.0330 | 1.000 |
| Dynamic range (IQR) | Effect of regressor (SD) | 16 | 0.532 | 0.0338 | 1.000 |
| Navigability | Effect of acquisition (SD) | 13 | -0.589 | 0.0343 | 1.000 |
| Mutated positions | Batch 10 − 100 at 400 assays | 16 | -0.493 | 0.0523 | 1.000 |
| Connected fraction | Batch 10 − 100 at 100 assays | 16 | 0.479 | 0.0604 | 1.000 |
| Connected fraction | Batch 10 − 100 at 400 assays | 16 | 0.465 | 0.0696 | 1.000 |
| Multi-mutant fraction | Effect of initial-set size (SD) | 16 | -0.459 | 0.0738 | 1.000 |
| Target density | Effect of representation (SD) | 16 | 0.459 | 0.0738 | 1.000 |
| Rank epistasis | Effect of initial-set size (SD) | 9 | -0.617 | 0.0769 | 1.000 |
| Connected fraction | Effect of batch size (SD) | 16 | -0.440 | 0.0882 | 1.000 |
| Connected fraction | Effect of acquisition (SD) | 16 | -0.433 | 0.0941 | 1.000 |
| Navigability | Greedy − EI at 400 assays | 13 | -0.470 | 0.1047 | 1.000 |
| Target density | Effect of initial-set size (SD) | 16 | 0.409 | 0.1159 | 1.000 |
| Local optima | Effect of acquisition (SD) | 13 | 0.445 | 0.1275 | 1.000 |
| Target density | Campaign difficulty at 400 assays | 16 | 0.362 | 0.1686 | 1.000 |
| Navigability | Batch 10 − 100 at 400 assays | 13 | 0.402 | 0.1737 | 1.000 |
| Target density | Effect of acquisition (SD) | 16 | 0.353 | 0.1800 | 1.000 |
| Rank epistasis | Effect of regressor (SD) | 9 | -0.483 | 0.1875 | 1.000 |
| Mutated positions | Campaign difficulty at 400 assays | 16 | -0.336 | 0.2039 | 1.000 |
| Assayed fraction | Effect of representation (SD) | 16 | 0.335 | 0.2043 | 1.000 |
| Library size | Effect of representation (SD) | 16 | -0.335 | 0.2043 | 1.000 |
| Assayed fraction | Effect of initial-set size (SD) | 16 | 0.332 | 0.2085 | 1.000 |
| Library size | Effect of initial-set size (SD) | 16 | -0.332 | 0.2085 | 1.000 |
| Mutated positions | Effect of initial-set size (SD) | 16 | 0.330 | 0.2124 | 1.000 |
| Navigability | Effect of batch size (SD) | 13 | -0.366 | 0.2189 | 1.000 |
| Local optima | Greedy − EI at 400 assays | 13 | 0.363 | 0.2233 | 1.000 |
| Mutated positions | Effect of representation (SD) | 16 | 0.315 | 0.2348 | 1.000 |

| | | | | | |
|---|---|---|---|---|---|
| Dynamic range (IQR) | Greedy − EI at 400 assays | 16 | 0.309 | 0.2445 | 1.000 |
| Dynamic range (IQR) | Batch 10 − 100 at 400 assays | 16 | -0.309 | 0.2445 | 1.000 |
| Connected fraction | Effect of regressor (SD) | 16 | 0.304 | 0.2524 | 1.000 |
| Dynamic range (IQR) | Effect of batch size (SD) | 16 | 0.297 | 0.2639 | 1.000 |
| Assayed fraction | Greedy − EI at 400 assays | 16 | -0.277 | 0.2999 | 1.000 |
| Library size | Greedy − EI at 400 assays | 16 | 0.277 | 0.2999 | 1.000 |
| Multi-mutant fraction | Greedy − EI at 100 assays | 16 | -0.274 | 0.3050 | 1.000 |
| Connected fraction | Effect of representation (SD) | 16 | 0.272 | 0.3086 | 1.000 |
| Local optima | Effect of initial-set size (SD) | 13 | -0.286 | 0.3440 | 1.000 |
| Navigability | Campaign difficulty at 400 assays | 13 | 0.283 | 0.3482 | 1.000 |
| Rank epistasis | Effect of representation (SD) | 9 | -0.350 | 0.3558 | 1.000 |
| Target density | Batch 10 − 100 at 400 assays | 16 | -0.247 | 0.3563 | 1.000 |
| Mutated positions | Greedy − EI at 400 assays | 16 | 0.241 | 0.3678 | 1.000 |
| Library size | Effect of regressor (SD) | 16 | 0.232 | 0.3865 | 1.000 |
| Assayed fraction | Effect of regressor (SD) | 16 | -0.232 | 0.3865 | 1.000 |
| Target density | Effect of batch size (SD) | 16 | 0.224 | 0.4053 | 1.000 |
| Rank epistasis | Effect of batch size (SD) | 9 | -0.300 | 0.4328 | 1.000 |
| Multi-mutant fraction | Batch 10 − 100 at 400 assays | 16 | 0.206 | 0.4443 | 1.000 |
| Multi-mutant fraction | Effect of batch size (SD) | 16 | -0.206 | 0.4443 | 1.000 |
| Multi-mutant fraction | Effect of regressor (SD) | 16 | -0.203 | 0.4510 | 1.000 |
| Mutated positions | Effect of acquisition (SD) | 16 | 0.193 | 0.4744 | 1.000 |
| Connected fraction | Greedy − EI at 100 assays | 16 | 0.192 | 0.4775 | 1.000 |
| Navigability | Effect of representation (SD) | 13 | -0.206 | 0.4988 | 1.000 |
| Connected fraction | Effect of initial-set size (SD) | 16 | -0.182 | 0.4990 | 1.000 |
| Multi-mutant fraction | Batch 10 − 100 at 100 assays | 16 | 0.182 | 0.4991 | 1.000 |
| Dynamic range (IQR) | Effect of initial-set size (SD) | 16 | 0.176 | 0.5133 | 1.000 |
| Target density | Batch 10 − 100 at 100 assays | 16 | -0.173 | 0.5204 | 1.000 |
| Assayed fraction | Batch 10 − 100 at 100 assays | 16 | -0.168 | 0.5349 | 1.000 |
| Library size | Batch 10 − 100 at 100 assays | 16 | 0.168 | 0.5349 | 1.000 |
| Local optima | Batch 10 − 100 at 400 assays | 13 | -0.187 | 0.5411 | 1.000 |
| Navigability | Effect of initial-set size (SD) | 13 | -0.184 | 0.5466 | 1.000 |
| Multi-mutant fraction | Effect of acquisition (SD) | 16 | -0.159 | 0.5569 | 1.000 |
| Navigability | Batch 10 − 100 at 100 assays | 13 | 0.179 | 0.5589 | 1.000 |
| Dynamic range (IQR) | Effect of acquisition (SD) | 16 | 0.156 | 0.5643 | 1.000 |
| Assayed fraction | Effect of acquisition (SD) | 16 | 0.153 | 0.5717 | 1.000 |
| Assayed fraction | Batch 10 − 100 at 400 assays | 16 | -0.153 | 0.5717 | 1.000 |

| | | | | | |
|---|---|---|---|---|---|
| Library size | Batch 10 − 100 at 400 assays | 16 | 0.153 | 0.5717 | 1.000 |
| Library size | Effect of acquisition (SD) | 16 | -0.153 | 0.5717 | 1.000 |
| Rank epistasis | Batch 10 − 100 at 400 assays | 9 | 0.217 | 0.5755 | 1.000 |
| Library size | Effect of batch size (SD) | 16 | -0.147 | 0.5868 | 1.000 |
| Assayed fraction | Effect of batch size (SD) | 16 | 0.147 | 0.5868 | 1.000 |
| Multi-mutant fraction | Greedy − EI at 400 assays | 16 | -0.147 | 0.5868 | 1.000 |
| Local optima | Effect of regressor (SD) | 13 | -0.165 | 0.5905 | 1.000 |
| Local optima | Greedy − EI at 100 assays | 13 | 0.138 | 0.6541 | 1.000 |
| Mutated positions | Greedy − EI at 100 assays | 16 | 0.116 | 0.6678 | 1.000 |
| Target density | Greedy − EI at 100 assays | 16 | 0.116 | 0.6681 | 1.000 |
| Local optima | Batch 10 − 100 at 100 assays | 13 | -0.126 | 0.6808 | 1.000 |
| Multi-mutant fraction | Campaign difficulty at 400 assays | 16 | 0.097 | 0.7207 | 1.000 |
| Rank epistasis | Greedy − EI at 400 assays | 9 | -0.133 | 0.7324 | 1.000 |
| Local optima | Effect of batch size (SD) | 13 | 0.104 | 0.7343 | 1.000 |
| Local optima | Campaign difficulty at 400 assays | 13 | -0.104 | 0.7343 | 1.000 |
| Rank epistasis | Effect of acquisition (SD) | 9 | -0.117 | 0.7650 | 1.000 |
| Dynamic range (IQR) | Greedy − EI at 100 assays | 16 | -0.080 | 0.7699 | 1.000 |
| Target density | Effect of regressor (SD) | 16 | -0.079 | 0.7700 | 1.000 |
| Dynamic range (IQR) | Batch 10 − 100 at 100 assays | 16 | -0.076 | 0.7783 | 1.000 |
| Connected fraction | Campaign difficulty at 400 assays | 16 | 0.071 | 0.7924 | 1.000 |
| Dynamic range (IQR) | Campaign difficulty at 400 assays | 16 | -0.065 | 0.8118 | 1.000 |
| Local optima | Effect of representation (SD) | 13 | -0.071 | 0.8166 | 1.000 |
| Rank epistasis | Campaign difficulty at 400 assays | 9 | 0.067 | 0.8647 | 1.000 |
| Connected fraction | Greedy − EI at 400 assays | 16 | -0.043 | 0.8746 | 1.000 |
| Mutated positions | Effect of regressor (SD) | 16 | -0.040 | 0.8838 | 1.000 |
| Navigability | Greedy − EI at 100 assays | 13 | -0.044 | 0.8863 | 1.000 |
| Rank epistasis | Batch 10 − 100 at 100 assays | 9 | -0.050 | 0.8984 | 1.000 |
| Target density | Greedy − EI at 400 assays | 16 | -0.024 | 0.9311 | 1.000 |
| Assayed fraction | Greedy − EI at 100 assays | 16 | -0.012 | 0.9655 | 1.000 |
| Library size | Greedy − EI at 100 assays | 16 | 0.012 | 0.9655 | 1.000 |
| Navigability | Effect of regressor (SD) | 13 | 0.008 | 0.9787 | 1.000 |
| Rank epistasis | Greedy − EI at 100 assays | 9 | 0.000 | 1.0000 | 1.000 |
| Library size | Campaign difficulty at 400 assays | 16 | 0.000 | 1.0000 | 1.000 |
| Assayed fraction | Campaign difficulty at 400 assays | 16 | 0.000 | 1.0000 | 1.000 |

#### Supporting Figures

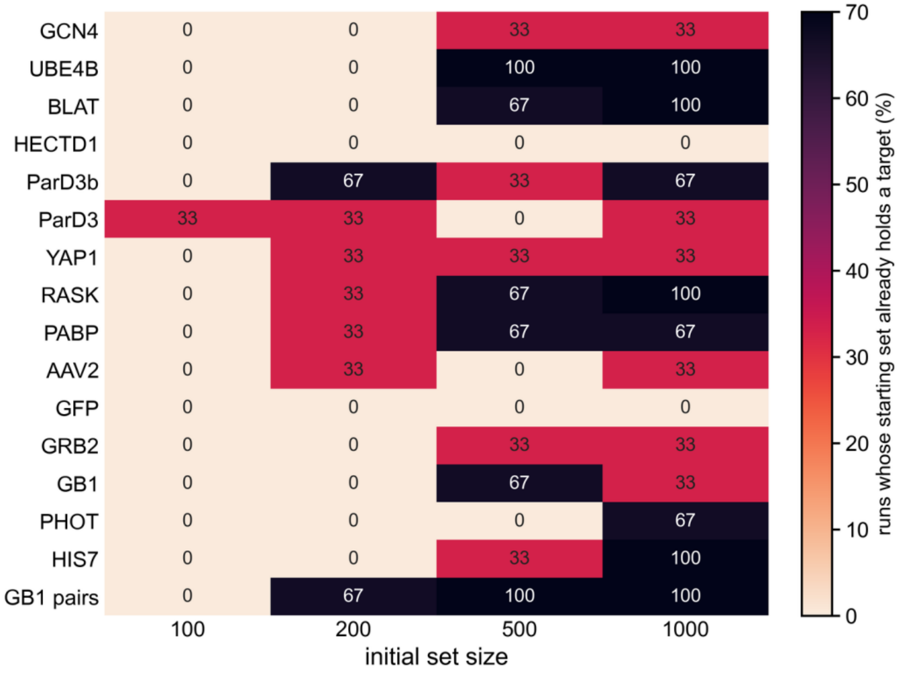


**Figure S1. Campaigns already successful at initialization.** Percentage of simulated campaigns whose initial training set already contained a variant in the highest-fitness 0.1%, shown for each landscape and initial training-set size. These campaigns were excluded from comparisons between MLDE configurations but retained in comparisons with random and zero-shot screening (**Method S4**).

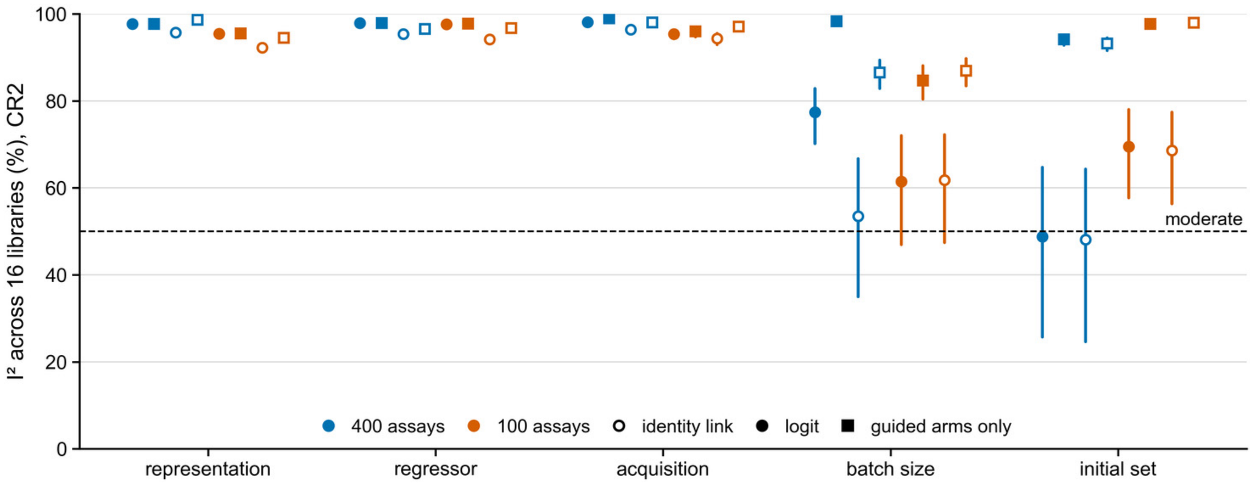


**Figure S2. Variation in the effects of MLDE design choices between protein fitness landscapes.** The $I^2$ statistic summarizes how much variation in the effect of each design choice reflects differences between landscapes rather than sampling uncertainty. Results are shown for all eight analysis specifications: two assay budgets (100 and 400 acquisitions), two link functions (logit and identity), and two campaign sets (all acquisition arms or ML-guided arms only). Estimates use CR2-corrected standard errors[7,8] and Higgins–Thompson confidence intervals.[6] The dashed line marks 50% $I^2$, a conventional reference point for moderate heterogeneity.[48] See **Method S5**.

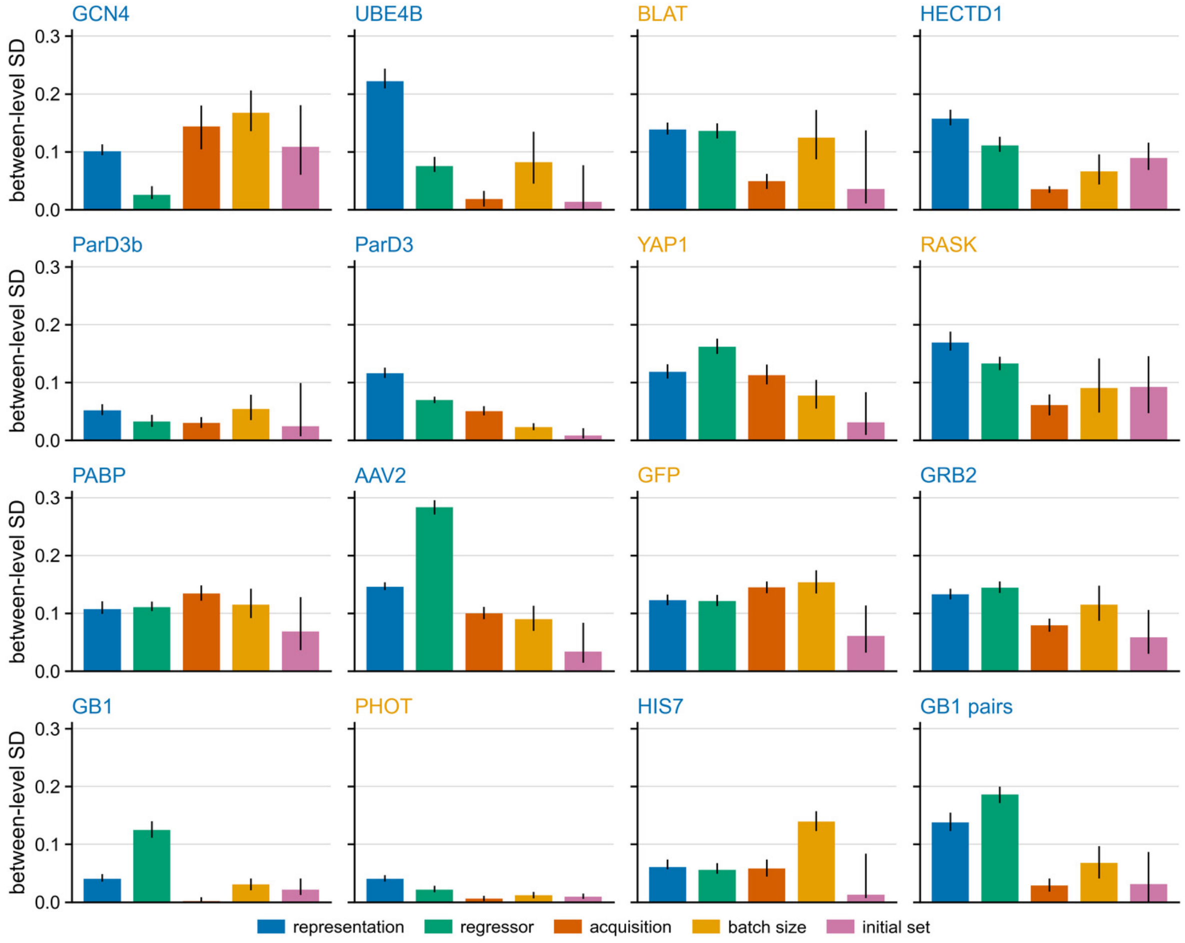


**Figure S3. Influence of individual MLDE design choices on campaign success in each landscape.** The effect of each design choice is quantified by the standard deviation of success rates across its tested levels after 400 acquired assays. Error bars indicate 95% confidence intervals from resampling complete scenarios. Landscape names are colored by library type.

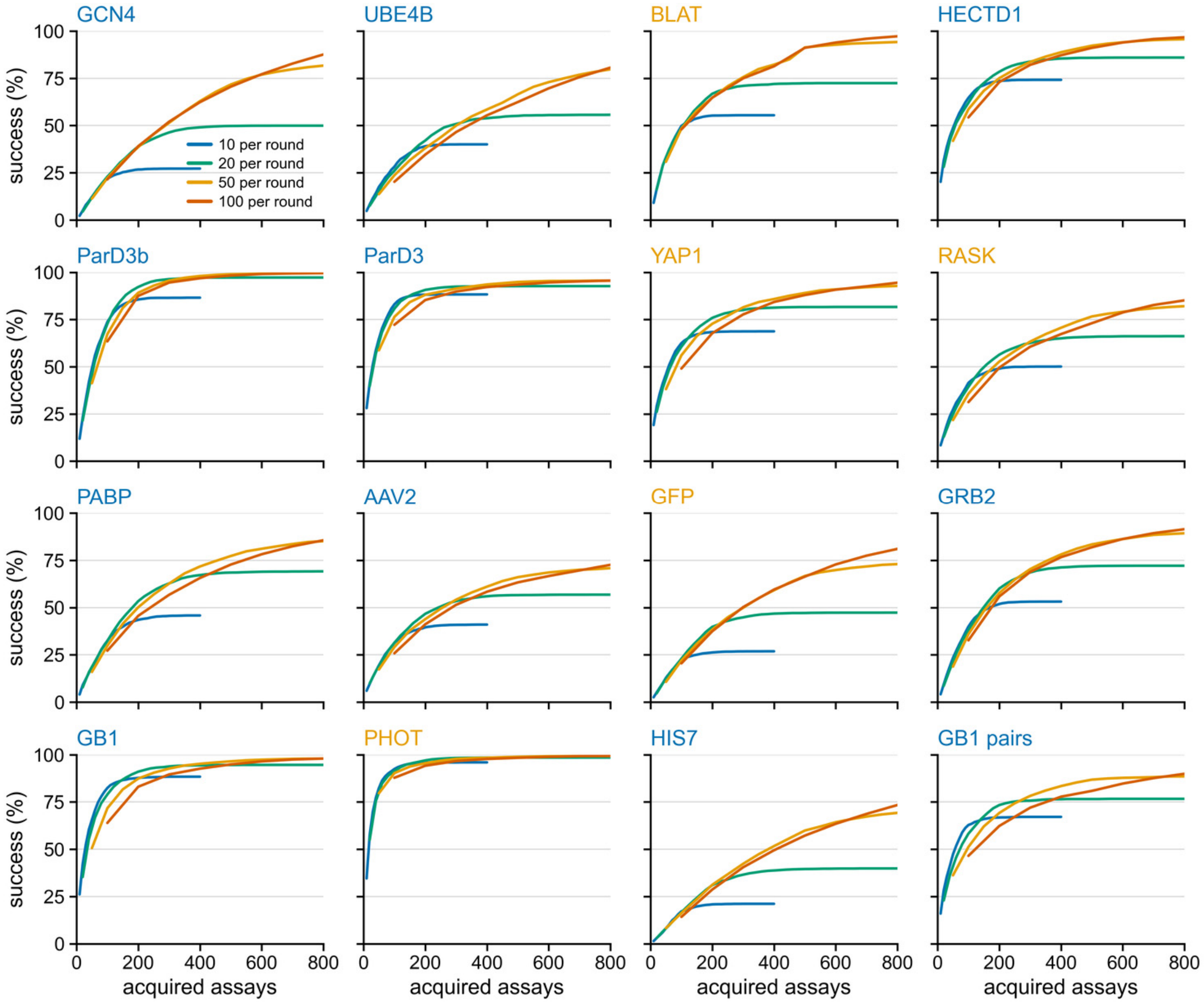


**Figure S4. Campaign success as a function of assay budget and acquisition batch size.** For each of the 16 landscapes, success rates are shown against the number of newly acquired assays for batch sizes of 10, 20, 50, and 100 variants. Curves end when campaigns reach the maximum of 40 acquisition rounds.

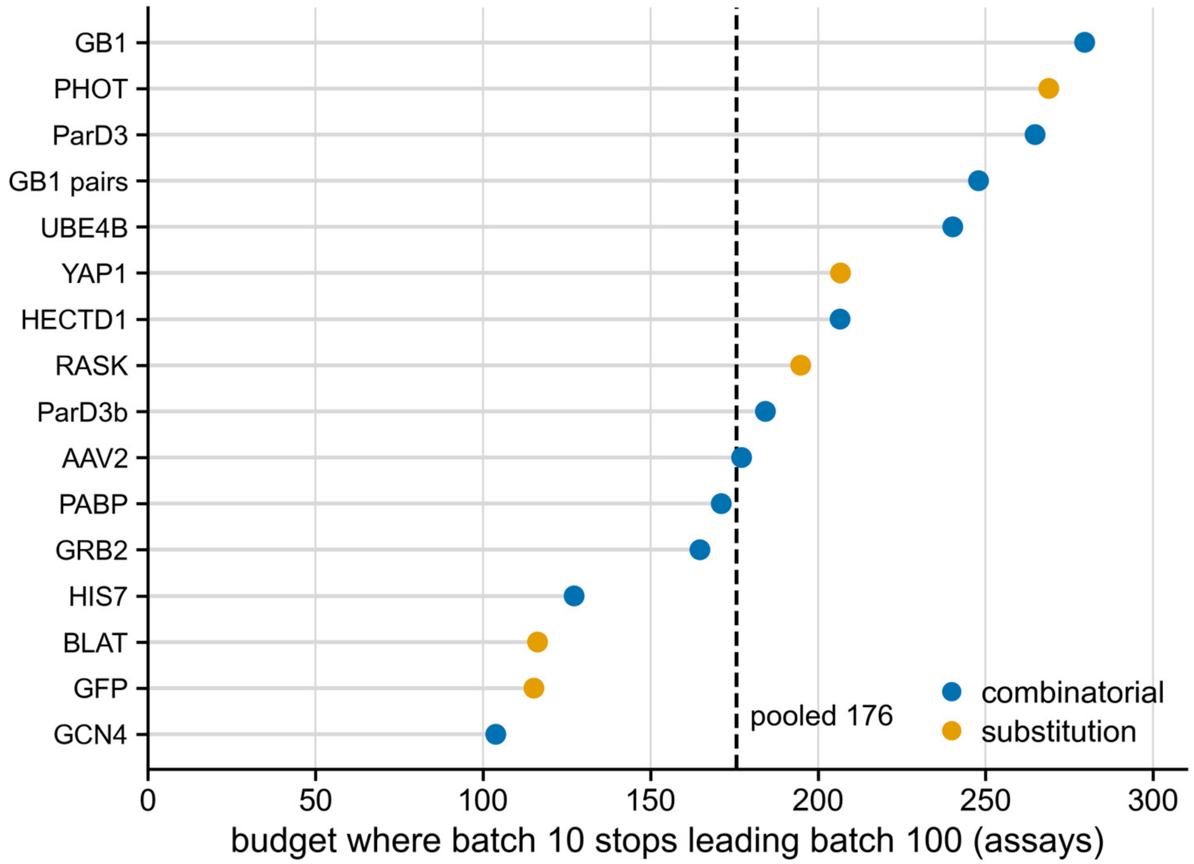


**Figure S5. Assay budget at which larger batches begin to outperform smaller batches.** For each landscape, the plot shows the number of newly acquired assays at which batches of 100 variants begin to achieve higher success rates than batches of 10. The dashed line marks the pooled crossover at 176 acquisitions. Because the main benchmark used a stopping rule defined by acquisition rounds, the crossover is partly influenced by that rule (**Method S3**; **Result S2**).

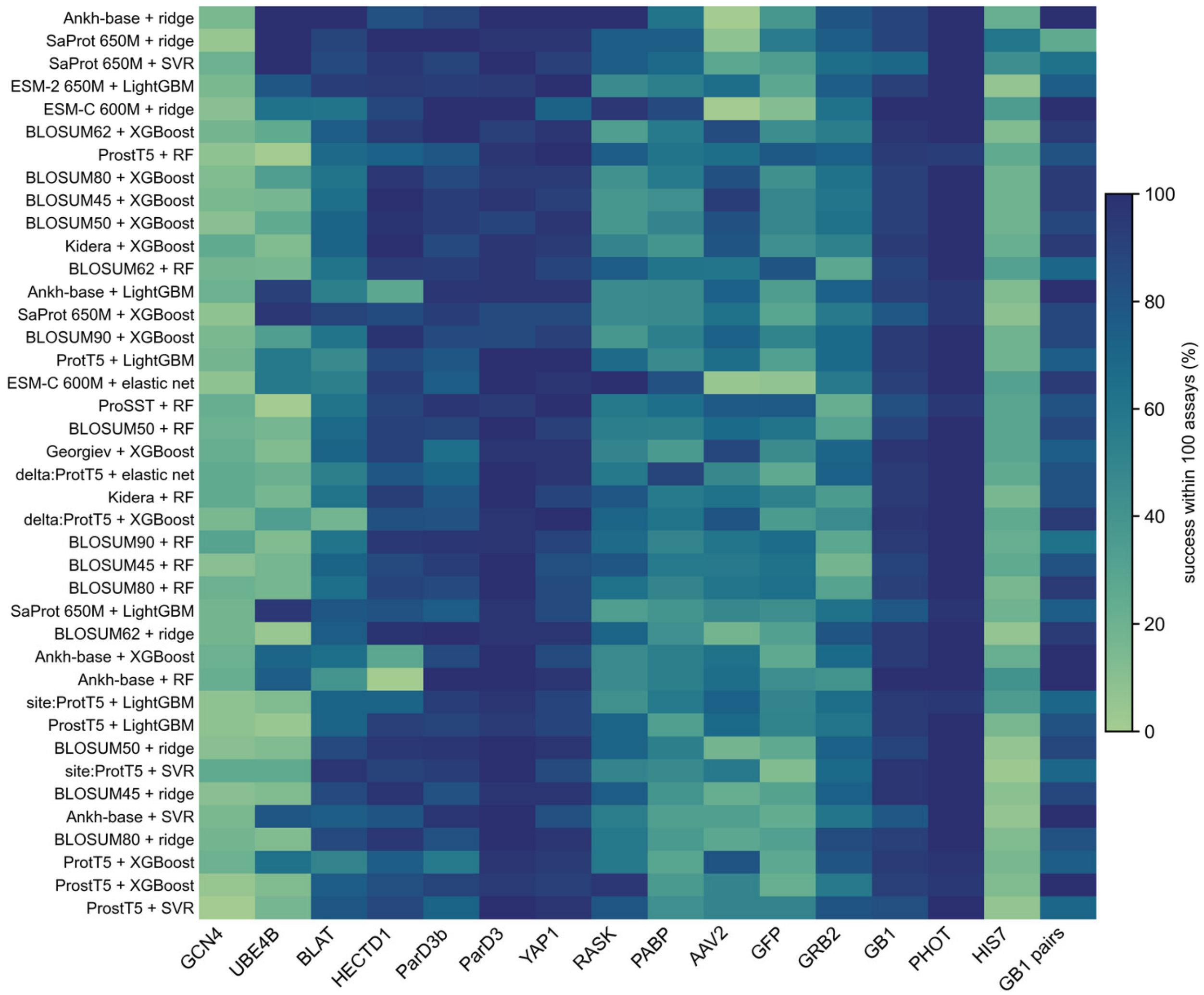


**Figure S6. Success rates of the 40 leading representation–regression-model combinations across landscapes.** The heatmap shows the success rate of each combination in each protein fitness landscape after 100 newly acquired assays using greedy acquisition. Landscapes are ordered by library size. **Tables S2** and **S4** list the representations and regression models.

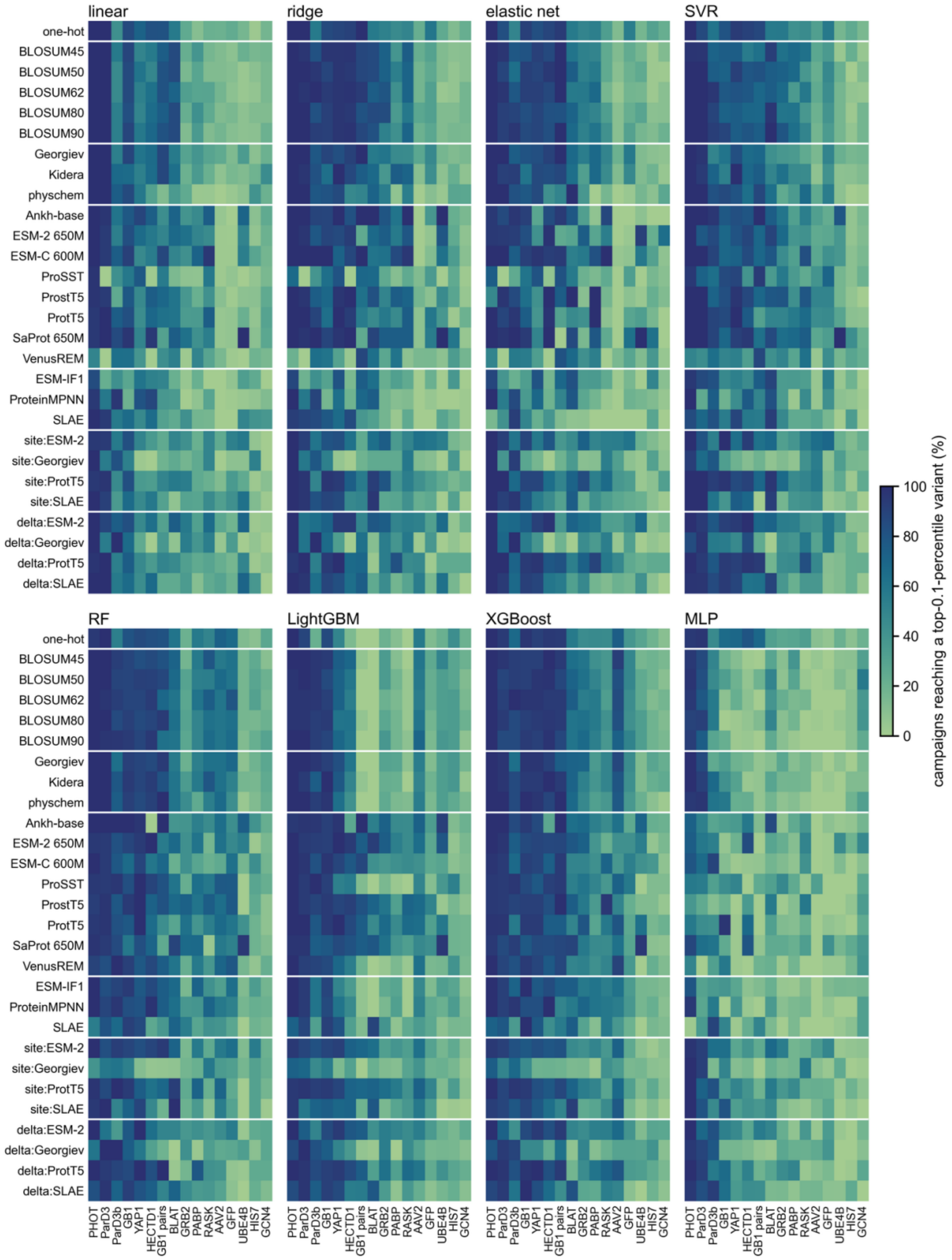


**Figure S7. Success rates of all representation–regression-model combinations after 100 acquired assays.** Each heatmap panel corresponds to one regression model. Rows show the 28 representations grouped into families and separated by white lines (one-hot; substitution matrices; physicochemical descriptors; protein language models; structure-based encoders; site-restricted variants; and variant-minus-wild-type [delta] representations). Columns show the 16 landscapes, ordered by average success across the grid. Color indicates the percentage of eligible campaigns that identified a top-0.1% variant using greedy acquisition. **Tables S2** and **S4** list the representations and regression models. The underlying values are provided in E6_all_combos_by_library.csv.

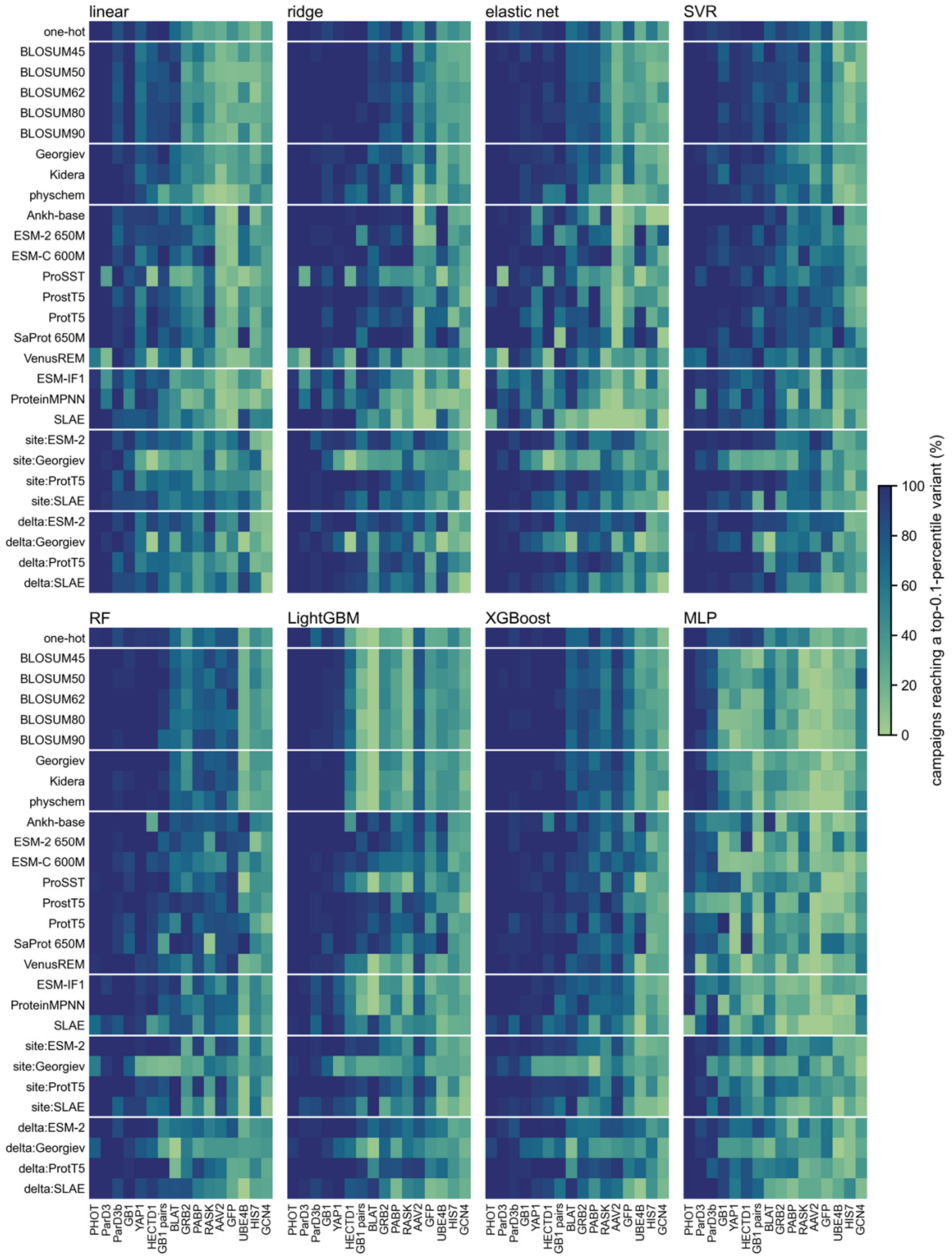


**Figure S8. Success rates of all representation–regression-model combinations after 200 acquired assays.** As in **Figure S7**, each panel shows one regression model, with the 28 representations grouped by family in rows and the 16 landscapes ordered by average success in columns. White lines separate representation families. Color indicates the percentage of eligible campaigns that identified a top-0.1% variant using greedy acquisition after 200 newly acquired assays. The underlying values are provided in E6_all_combos_by_library.csv.

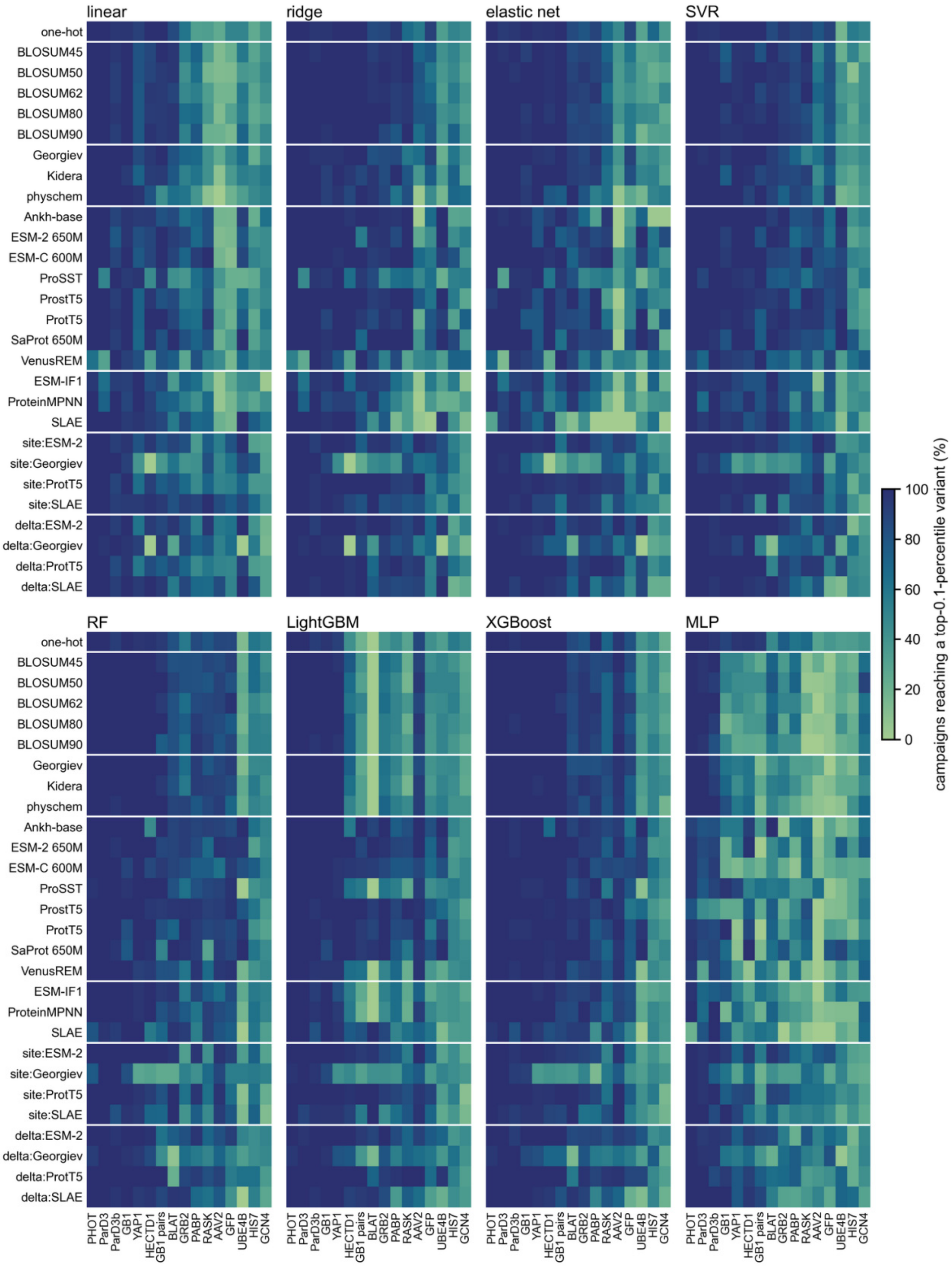


**Figure S9. Success rates of all representation–regression-model combinations after 400 acquired assays.** As in **Figures S7** and **S8**, each panel shows one regression model, with representations grouped by family in rows and the 16 landscapes ordered by average success in columns. White lines separate representation families. Color indicates the percentage of eligible campaigns that identified a top-0.1% variant using greedy acquisition after 400 newly acquired assays. The underlying values are provided in E6_all_combos_by_library.csv.

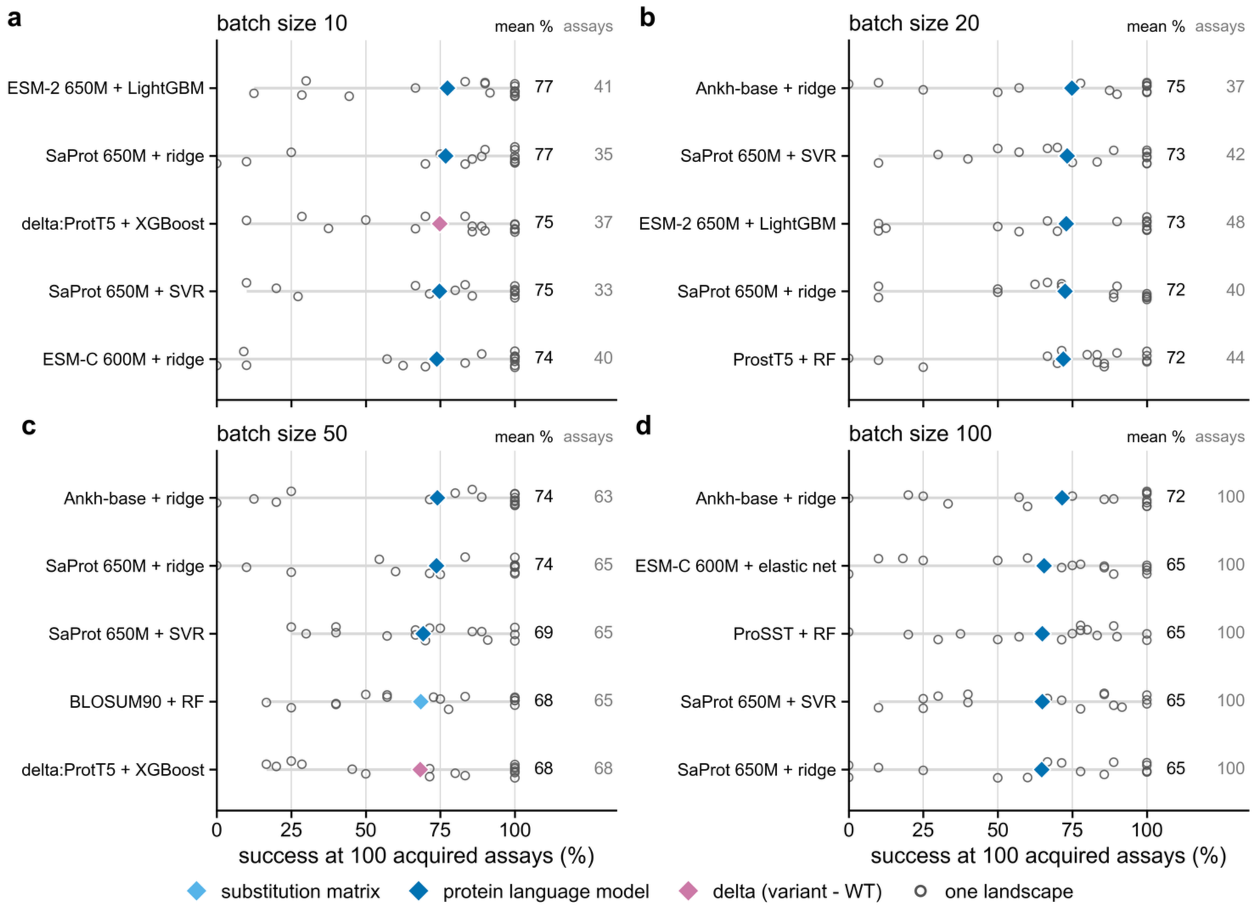


**Figure S10. Leading representation–regression-model combinations for each acquisition batch size after 100 acquired assays.** The analysis follows main-text **Figure 10** but evaluates success after 100 rather than 400 newly acquired assays. (a–d) The five combinations with the highest landscape-balanced success were ranked separately for each batch size and averaged across the four initial training-set sizes. Open circles indicate success rates in each of the 16 landscapes, gray lines show their range, and diamonds indicate landscape-balanced means, colored by representation family. The columns on the right report the mean success rate and the mean number of assays required to identify the first successful variant. Rankings were more stable than at 400 assays: 10 distinct combinations occupied the 20 top-five positions, compared with 17 at 400 assays. Nevertheless, half of the 20 leading entries, including three of the four top-ranked combinations, had at least one landscape in which no campaign identified a target. With a batch size of 100, the 100-assay budget allows only one acquisition round, so every successful campaign found its target at 100 acquired assays.

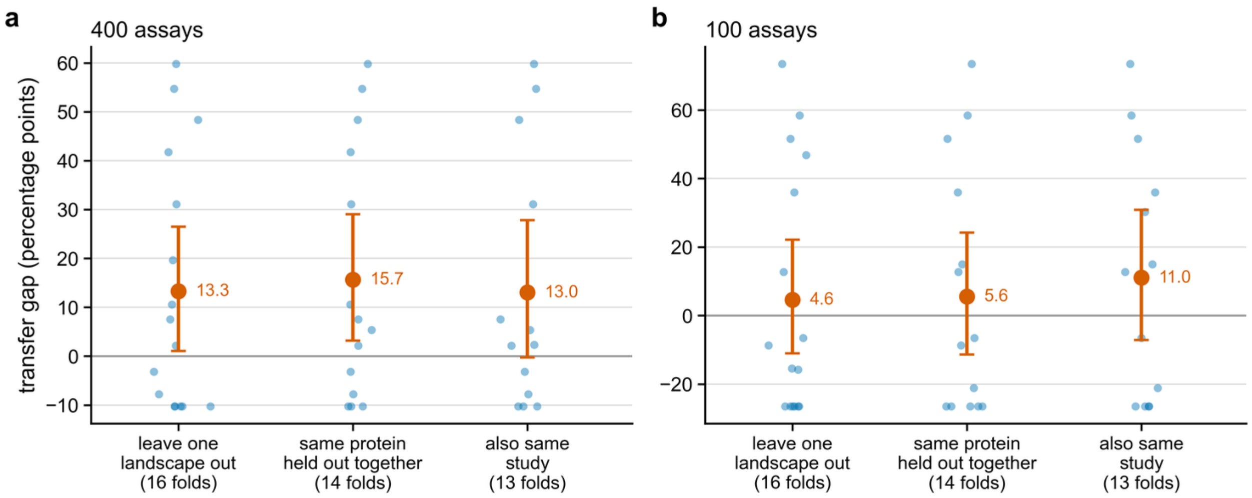


**Figure S11. Reduction in configuration performance on held-out landscapes under different exclusion schemes.** Results are shown after (a) 400 and (b) 100 newly acquired assays. Each point represents one held-out evaluation; larger markers show the mean performance reduction and error bars indicate 95% confidence intervals from bootstrapping landscapes. Under the standard leave-one-landscape-out scheme, each of the 16 landscapes was withheld in turn (16 folds). Under the same-protein scheme, landscapes from the same protein were withheld together (GB1 with GB1 pairs, SPG1_STRSG; ParD3 with ParD3b, F7YBW8_MESOW), giving 14 folds.[17,18,19,20] Under the same-study scheme, UBE4B and HECTD1, which came from the same study[26] and represent different proteins, were additionally withheld together, giving 13 folds. The reduction did not become smaller with either stricter scheme, suggesting that the estimate from the standard analysis was not explained by overlap between related landscapes. **Table S1** gives the dataset accessions.
